# Estimating the fitness cost and benefit of antimicrobial resistance from pathogen genomic data

**DOI:** 10.1101/2022.12.02.518824

**Authors:** David Helekal, Matt Keeling, Yonatan H Grad, Xavier Didelot

**Affiliations:** Centre for Doctoral Training in Mathematics for Real-World Systems, University of Warwick, UK; Mathematics Institute and School of Life Sciences, University of Warwick, UK; Department of Immunology and Infectious Diseases, TH Chan School of Public Health, Harvard University, USA; School of Life Sciences and Department of Statistics, University of Warwick, UK

**Keywords:** genomic epidemiology, phylodynamics, antimicrobial resistance, resistance fitness cost

## Abstract

Increasing levels of antibiotic resistance in many bacterial pathogen populations is a major threat to public health. Resistance to an antibiotic provides a fitness benefit when the bacteria is exposed to this antibiotic, but resistance also often comes at a cost to the resistant pathogen relative to susceptible counterparts. We lack a good understanding of these benefits and costs of resistance for many bacterial pathogens and antibiotics, but estimating them could lead to better use of antibiotics in a way that reduces or prevents the spread of resistance. Here, we propose a new model for the joint epidemiology of susceptible and resistant variants, which includes explicit parameters for the cost and benefit of resistance. We show how Bayesian inference can be performed under this model using phylogenetic data from susceptible and resistant lineages and that by combining data from both we are able to disentangle and estimate the resistance cost and benefit parameters separately. We applied our inferential methodology to several simulated datasets to demonstrate good scalability and accuracy. We analysed a dataset of *Neisseria gonorrhoeae* genomes collected between 2000 and 2013 in the USA. We found that two unrelated lineages resistant to fluoroquinolones shared similar epidemic dynamics and resistance parameters. Fluoroquinolones were abandoned for the treatment of gonorrhoea due to increasing levels of resistance, but our results suggest that they could be used to treat a minority of around 10% of cases without causing resistance to grow again.

## INTRODUCTION

The levels of antimicrobial resistance of many pathogens have risen worryingly over the past few decades. In a report on the threat posed by antibiotic resistance published by the CDC (Centres for Disease Control and Protection), three microorganisms including *N. gonorrhoeae* are classified as posing an urgent threat level, and twelve more represent a serious threat to public health [1]. A review on antimicrobial resistance estimated that resistance claims at least 700,000 lives per year worldwide and that the death toll could go up to 10 million per year by 2050 if current trends are allowed to continue [2], and a recent study estimated that there were almost 5 million deaths associated with resistance in 2019 [3]. Few new antimicrobials have been developed and deployed since the 1970s, whereas resistance to new drugs often emerges soon after initial introduction [4], so that several pathogens are dangerously close to becoming completely untreatable. Effectively tackling antimicrobial resistance requires greater understanding of epidemiological and evolutionary factors leading to emergence of resistance and the spread of resistance through pathogen populations. Achieving this goal requires development of mathematical models of antimicrobial resistance and robust statistical analysis of epidemiological models with informative observations. This modelling approach to resistance was initiated in the late 1990s [5, 6] and has led to the development of many models, appropriate for different organisms, mode of spread, study scale and context [7].

Resistance brings a clear fitness benefit to pathogens acquiring it in the presence of antimicrobials. The net value of this fitness benefit therefore increases with the frequency with which the specific antimicrobial is employed, either against the pathogen itself or more generally in the case of a pathogen that can be carried asymptomatically. However, resistance also typically comes with a fitness cost to the pathogen [8]. The simplest demonstration of this effect is when discontinued use of an antimicrobial leads to reductions in resistance rates. The fitness costs and benefits of resistance remain poorly understood for many pathogens and antimicrobials [9]. A better quantification of resistance benefits and costs is required to provide a solid basis for evaluating the potential effectiveness of public health intervention measures proposed to exploit fitness costs in the hope of stopping or even reversing the spread of resistance [9]. For example, the numbers of gonorrhoea cases sensitive and resistant to cefixime in England over a decade was recently analysed to quantify the cost and benefit associated with resistance to this antibiotic [10]. These estimates were used to predict that cefixime could be reintroduced to treat a minority (∼25%) of gonorrhoea cases without causing an increase in cefixime resistance levels, which would reduce the risk of emergence of resistance to the currently used antibiotics. Moreover, the extent of the fitness cost of resistance can vary by genomic background [11], such that the effect of interventions that seek to capitalize on the fitness costs of resistance may be lineage dependent. Therefore, it is necessary to estimate fitness costs at the per lineage level. The aim of this study is to quantify the contribution that changes in prescription policy have on the population dynamics of particular resistant lineages. This is in contrast to studies that are interested in the overall ecology of resistance or the eventual fate of a resistant phenotypes, see for example [12].

Pathogen genomic data has great potential to help us understand the evolutionary and epidemiological dynamics of infectious disease [13]. An important advantage of this phylodynamic approach is that analysis of genomic data is less sensitive to sampling biases, especially when using a coalescent framework which describes the ancestry process conditional on sampling [14]. A few studies have used this approach to shed light on the fitness cost associated with antimicrobial resistance. For example, a study showed the association between the growth rate of a methicillin-resistant *Staphylococcus aureus* lineage and consumption of beta-lactams [15]. Other studies quantified the relative transmission fitness of resistance mutations in HIV [16] and *Mycobacterium tuberculosis* [17]. Here, we take a different approach by modelling explicitly the phylodynamic trajectories of the sensitive and resistant lineages as a function of the fitness cost, which is constant, and the fitness benefit, which depends on the antimicrobial consumption. Our method therefore requires three inputs: the amount of antimicrobial being used over time, genomic data from a sensitive lineage, and genomic data from a resistant lineage. From this we disentangle the fitness cost and benefit of resistance, thereby providing the parameters needed to predict phylodynamic trajectories and inform recommendations on how to use antimicrobials without worsening the resistance threat. Overall, the scenario we are interested in is that of overall resistance dynamics at a large population level. In such a scenario the bulk of incidence is going to be caused by local transmission rather than imports. We do not intend for the methods presented in this paper to be applicable to small population dominated by imports and complex, heterogeneous routes of transmission, such as within hospital setting of hospital acquired infections. For such a scenario, a different approach using Birth-Death type models would be more appropriate [16, 17].

## METHODS

### Overall approach

Pathogen phylogenetic data contains information about past population size dynamics of the pathogen under study [13, 18]. Under assumptions of the epidemic process being characterised well enough by a simple compartmental epidemic model, this information about population size dynamics can be translated into epidemic trajectories [19, 20]. These epidemic trajectories can be described using an epidemic model which accounts for the effects of a fitness cost and benefit of resistance to a specific antimicrobial. As the use of this antimicrobial changes through time, so will the net fitness of the particular lineage in consideration. This will in turn lead to changes in the behaviour of the epidemic trajectory. However, not all changes in the behaviour of the epidemic trajectory will be due to changes in the fitness of the resistant phenotype. Confounding factors, such as as depletion of susceptibles or changes in host behaviour will also affect the epidemic trajectory. Under relatively mild assumptions detailed below changes in these confounding factors will affect other lineages equally. We can therefore use as “control” some data from a susceptible lineage, ideally closely related and with the same resistance profile to other antimicrobials used in significant amounts as primary treatment. Differences between the trajectories of the sensitive and resistant lineages can then be ascribed specifically to resistance, allowing us to estimate the associated fitness cost and benefit parameters.

Let us consider a pathogen causing infections at the level of a large population that are or were treated with a certain antimicrobial compound. We assume that at some point in the past one or several lineages with resistance to this antimicrobial compound have arisen. Our aim is to quantify the fitness cost and benefit of the resistance to this antimicrobial for a given lineage as a function of use of the antimicrobial of interest through time. To this end we need data that quantify the use over time of the given antimicrobial to treat infections caused by this pathogen, as well as a reasonable sample of sequenced case isolates from infections caused by the pathogen over time. Furthermore, we need information that characterises the resistance profiles of the individual isolates, which can be obtained either by resistance screening *in vitro*, or predicted from the sequences *in silico* [21]. A dated phylogeny of these samples is estimated, for example using BEAST [22], BEAST2 [23] or BactDating [24]. This phylogeny is then used as the starting point for analysis [25], to identify which samples belong to resistant and susceptible lineages and to select related lineages for further study that are wholly resistant or susceptible to the antimicrobial of interest, but otherwise similar in their resistance profiles. Note that for simplicity resistance is treated as a binary trait, with samples being either resistant or susceptible to antimicrobials, as is usually the case in resistance modelling studies [7].

### Transmission model derivation

In order to estimate the fitness cost and benefit of antimicrobial resistance, a transmission model needs to be specified. We focus on estimating the fitness parameters of a particular lineage harbouring a certain treatment resistant phenotype when previous infection does not confer immunity against reinfection. Under the simplifying assumptions that the host population is unstructured and that past infections do not confer any immunity, the multi-lineage Susceptible-Infected-Susceptible (SIS) is a reasonable model [26, 27]. This model is more commonly referred to as multi-strain SIS. Fluctuations in the carriage levels of different lineages can also be due to external factors, such as changes in host demography or behaviours. Left unaccounted, such fluctuations would bias estimates of the fitness cost and benefit of resistance to a given antimicrobial. Therefore, we modify the model with time-varying transmission rate *β*(*t*) and population size *N*(*t*). This leads to an *n*-lineage model described by a system of the following *n*-coupled ordinary differential equations (ODEs):

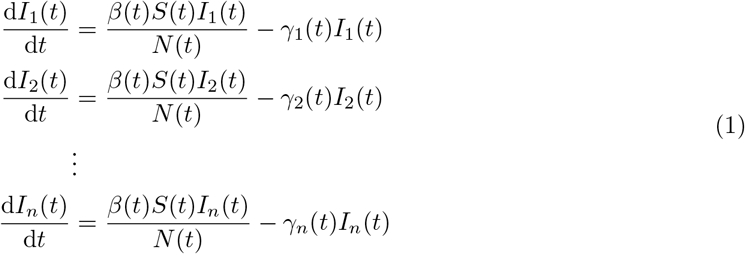

Where *I*_*j*_(*t*) denotes the number of people infected with the *j*-th lineage at time *t. β*(*t*) is the transmission rate that varies with time due for example to changes that are not specific to any lineage, for example host behaviour. *N* (*t*) is the host population size which may also change with time due to demographic factors. *γ*_*j*_(*t*) is the recovery rate of the *j*-th lineage at time *t*. These may or may not vary with time through their dependency on the antimicrobial usage which changes with time. Finally *S*(*t*) denotes the number of susceptible hosts

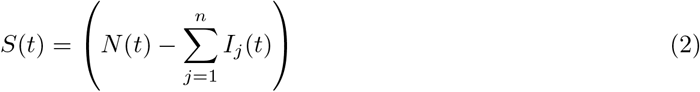

Typically this model could simply be reduced to a two lineage model, averaging over all lineages that are phenotypically similar in their resistance profiles. However, this is undesirable, as some of the lineages with the same resistance phenotype could differ in fitness due to different genomic background which would confound our estimates. Furthermore this sort of model would not be readily tractable in a genomic framework, because phylogenetic data is generally going to be informative about the dynamics of a particular lineage only. Note that this also means that the analysis produced is valid for the lineages being studied, and cannot be extrapolated to the overall dynamics of resistance for a given pathogen.

We therefore need to focus on the resolution of individual lineages. We note that environmental effects such as fluctuations in host population size or behaviour affect all lineages equally, if the population is well mixed. We denote the combination of these effects as *b*(*t*) = *β*(*t*)*S*(*t*)/*N* (*t*). Conditional on the knowledge trajectory of *b*(*t*) the ODEs in Equation 1 become uncoupled, and this allows us to reduce the system to uncoupled equations corresponding to the lineage we will be focusing on. As such we will treat *b*(*t*) as a random object that needs to be inferred. We further assume that for the susceptible lineages the average recovery rate denoted *γ*_*s*_ does not change over time, whereas for the resistant lineage it takes one of two values: *γ*_*T*_ = *q*_*T*_ + *γ*_*s*_ if a given patient is treated with the antimicrobial of interest, or *γ*_*U*_ = *q*_*U*_ + *γ*_*s*_ otherwise. If we also consider the known proportion of registered cases treated with the antimicrobial of interest *u*(*t*), this fully determines the average recovery rate of the resistant lineages as:

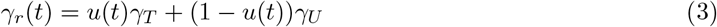

We can now fully write down the equations of the model we will be using for the sensitive and resistant lineages, respectively:

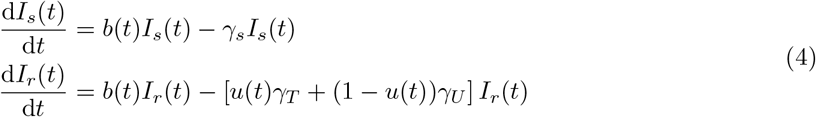

In practice, we are interested in the difference in recovery rates between the susceptible and the resistant lineage and sensitive lineage when every case gets treated with the antimicrobial of interest, and when the antimicrobial of interest is not used at all. We denote these by

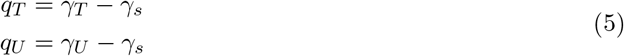

The interpretation is therefore that *q*_*T*_ captures the fitness benefit of resistance in the case *q*_*T*_ < 0 and *q*_*U*_ captures the fitness cost of resistance in the case *q*_*U*_ > 0.

This model can be applied to any number of resistant and sensitive lineages, simply by adding lineages associated terms to the likelihood and adding required parameters. This is straightforward as the individual lineages are independent conditional on *b*(*t*). but for simplicity the remainder of methods description focuses on the case of a single sensitive and a single resistant lineage, with the general case being a straightforward extension.

### Link to phylogenies

Having defined the epidemiological model, we can now link it to the phylogenetic process. Based on [19, 28], the instantaneous coalescent rates for a single pair of lineages can be derived as

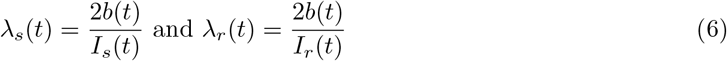

in the susceptible and resistant populations, respectively. The likelihood of a dated phylogeny **g** with *n* leaves at times *s*_1_ < 舰 < *s*_*n*_ and *n* −1 coalescent events at times *c*_1_ < 舰 < *c*_*n*−1_ and *A*(*t*) lineages at time *t* is therefore given by [29]:

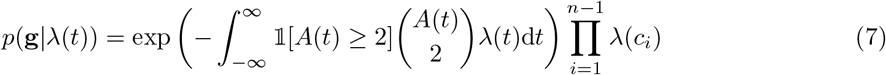

Where *λ*(*t*) = *λ*_*s*_(*t*) and *λ*(*t*) = *λ*_*r*_(*t*) for the susceptible and resistant phylogenies, respectively. However, in most cases, and indeed in our case, the integral in Equation 7 is not analytically intractable. Furthermore, the antibiotic use data is unlikely to span the entire phylogeny. Therefore, we define the approximate likelihood for the phylogeny truncated to [*t*_min_, *t*_max_], which is the intersection interval spanned by the antibiotic use data and the phylogenies under study.

As such we resort to the standard way of approximating coalescent likelihoods [30], partitioning the interval [*t*_min_, *t*_max_] into a fine mesh *t*_min_ = *t*_1_ < *t*_2_ < *t*_3_ < 舰 < *t*_*N*_ = *t*_max_ such that *t*_*i*_ − *t*_*i*−1_ < Δ_*t*_ and that all sampling and coalescent times between *t*_min_ and *t*_max_ are included in the mesh:

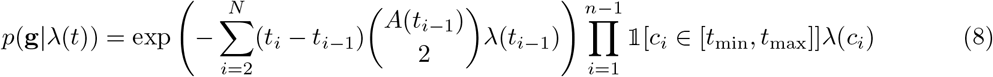

We note that the approach of how we treat the relationship between the phylogenies and epidemic is effectively a structured coalescent with no migration and time varying *N*_e_(*t*) determined by the deterministic epidemic model. Approaches reminiscent of ours have been used to formally study the expected age of a mutation in both presence or absence of selection [31]. However in that case the populations correspond to different alleles, and the *N*_e_(*t*) curves follow the proportion of population with a given allele as determined by Wright-Fisher diffusion forwards in time. Migration between the demes corresponding to individual alleles can also further be added corresponding to recombination [32].

### Bayesian inference

We first re-scale time from the interval [*t*_min_, *t*_max_] to [−1, 1]. Denoting the scale factor *D* = (*t*_max_ −*t*_min_)/2 associated with this re-scaling, we account for this in the model by defining 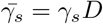. The model consists of independent first-order linear homogeneous ODEs for each lineage with time-varying coefficients. The solutions at time *t* subject to initial conditions *I*_*s*_(0) = *I*_*s*0_ and *I*_*r*_(0) = *I*_*r*0_ can be obtained in terms of the integral of the instantaneous rates up to time *t*:

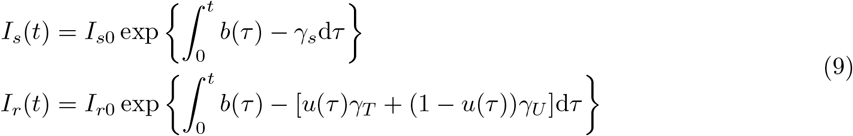

As it stands, this model would not be well-suited for performing inference under, primarily due to the difficulty in choosing a sensible prior on *b*(*t*), and a very complicated dependency structure between the initial conditions and *b*(*t*). As such we re-parameterise the model by directly modelling the logarithm of *I*_*s*_(*t*) as a Gaussian Process:

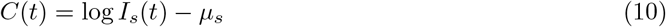

Where *C*(*t*) is an appropriately chosen zero mean Gaussian Process, and *μ*_*s*_ is the susceptible intercept which relates to the susceptible initial condition *I*_*s*0_ as follows:

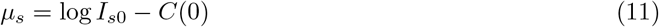

We use this formulation principally to loosen the coupling between the intercept parameter and the Gaussian Process in order to speed up sampling. From this we can compute *b*(*t*) and log *I*_*r*_(*t*) as

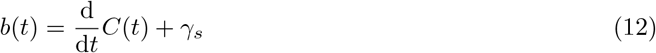

and

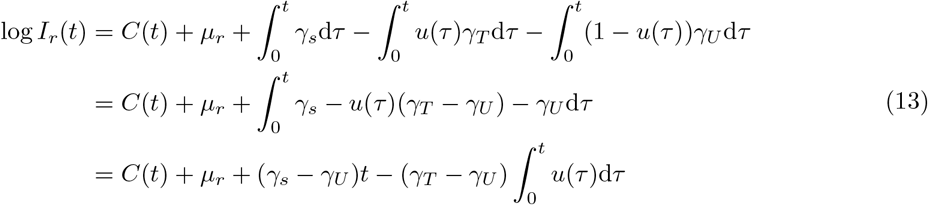

Once again we follow the same reasoning for the resistant trajectory intercept *μ*_*r*_, relating it to *I*_*r*0_ as:

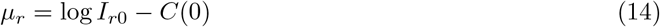

Note that 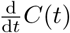 exists as long as the associated covariance kernel is sufficiently smooth such as in the case of the radial basis function (RBF) kernel [33] which we used. Evaluating a full-rank, Gaussian process with differentiable trajectories on the entirety of the mesh would be prohibitively expensive due to the *O*(*n*^3^) computational complexity, where *n* is the number of grid points. Such a high computational cost would make the model infeasible. Instead, we work with a low-rank representation of *C*(*t*) based on the framework introduced in [34]. This leads to the representation of the low-rank projection of *C*(*t*), denoted by *Ĉ*(*t*)

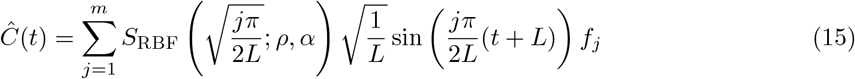

and

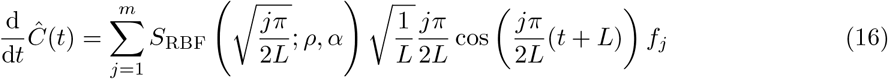

This reduces the evaluation complexity of the Gaussian process prior from *O*(*n*^3^) to *O*(*nm*). *L* and *m* are approximation parameters that need to be specified a-priori, see [34] for details. In practice we used the Hilbert Space Gaussian Process (HSGP) approximation with parameters *L* = 6.5 and *m* = 60. These approximation parameters are appropriate for the 99% interval of the length-scale prior used as per [34]. Where *f*_*j*_ are independent and identically distributed random variables following the standard Gaussian distribution, *S*_RBF_(·; ·,·) is the appropriate spectral density for the RBF kernel, *ρ* is the kernel length scale and *α* is the marginal standard deviation of the kernel [34].

Denote by ***θ*** = (*γ*_*s*_, *γ*_*U*_, *γ*_*T*_, *I*_*s*0_, *I*_*r*0_, *Ĉ*(*t*)) the parameters of the pathogen dynamics model. We can now factorise the model posterior *π*(***θ***, *α, ρ, f*_1:*m*_ | **g**_*s*_, **g**_*r*_), suppressing dependency on *t* where appropriate:

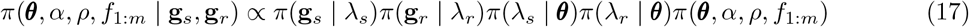

The first two terms are computed using the coalescent likelihood in Equation 7. The third term is given by combining Equations 6, 10 and 12. The fourth term is obtained by combining Equations 6, 12 and 13. Finally, the last term is given by:

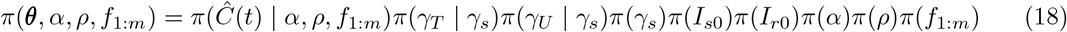

where the first term is given by the Gaussian process (Equations 15 and 16) and the remaining terms correspond to the prior distributions listed below.

### Choice of prior and parameterisation

The model is parameterised with the priors summarised in Table 1. The data is not expected to be very informative about the value of *γ*_*s*_. As such, we impose a fairly informative prior on this parameter, centred around a guess *γ*^*^ which must be known and supplied *a priori. σ* then governs how informative the prior is. We typically use a value of *σ* = 0.3, which includes relative fluctuations of over 50% in its 95% interval. The higher the value of *σ*, the more complicated the geometry and subsequently sampling of the posterior becomes. *γ*_*T*_ and *γ*_*U*_ represent the recovery rates for the resistant lineage when the resistant lineage is treated with the focal antibiotic of interest, or another antibiotic, respectively. A normal distribution centred at *γ*_*s*_ and truncated to positive values only is a natural choice. We choose its standard deviation to be 0.3*γ*^*^ as this puts > 99% of the weight within 2*γ*^*^ thus making implausibly large fluctuations unlikely. Such large fluctuations are hardly of interest here since they would lead to a very rapid selective sweep or extinction. The recovery rates *γ*_*T*_ and *γ*_*U*_ are related to the absolute changes in recovery and therefore fitness parameters using Equation 5. *γ*_*U*_ *> γ*_*s*_ corresponds to faster recovery when the resistant lineage is treated with an antimicrobial it is sensitive to and therefore a cost of resistance. *γ*_*T*_ < *γ*_*s*_ corresponds to slower recovery when the resistant lineage is treated with the antimicrobial of interest and therefore a benefit of resistance. If instead a large proportion of posterior probability mass has *γ*_*U*_ < *γ*_*s*_ or *γ*_*T*_ > *γ*_*s*_, we conclude that the result is consistent with either the cost or the benefit of resistance not being significantly present. The prior on *ρ* was chosen so that approximately 1% of mass lies on values of *ρ* < 0.2 and approximately 1% of mass lies on *ρ* > 2. The lower bound was chosen to avoid over-fitting, and the upper bound to suppress length scales that exceed the range of data and thus cannot be informed about by the data.

**Table 1:** Summary of the parameters and priors used in the model.

| Parameter | Symbol | Prior |
| --- | --- | --- |
| Susceptible lineage recovery rate | $\gamma_s$ | `log-normal`($\log \gamma^*, \sigma$) |
| Resistant lineage recovery rate if treated with focal antibiotic | $\gamma_T$ | `normal`($\gamma_s, 0.3\gamma^*$)$\mathbb{1}[x > 0]$ |
| Resistant lineage recovery rate if treated with other antibiotic | $\gamma_U$ | `normal`($\gamma_s, 0.3\gamma^*$)$\mathbb{1}[x > 0]$ |
| Initial prevalence of sensitive lineage | $I_{s0}$ | `log-normal`(6, 2) |
| Initial prevalence of resistant lineage | $I_{r0}$ | `log-normal`(6, 2) |
| GP kernel marginal variance | $\alpha$ | `gamma`(4, 4) |
| GP kernel length scale | $\rho$ | `inverse-gamma`(4.63, 2.21) |
| Approximate GP functions | $f_{1:m}$ | $\mathcal{N}(0, 1)$ |

In practice, due to our choice of a sampling approach we need to parametrise *γ*_*U*_ and *γ*_*T*_ on an unconstrained space, and ideally also weaken the dependency on *γ*_*s*_. To do so we introduce parameters 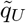 and 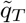, and define *γ*_*U*_ and *γ*_*T*_ to be a deterministic transformation of these

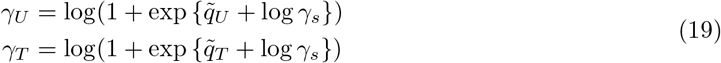

The Jacobian adjustment to the likelihood associated with this transformation is proportional to

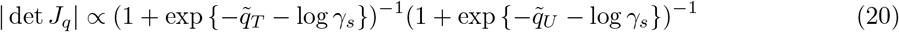

### Computational implementation

The posterior in Equation 17 is a high dimensional distribution and we expect many parameters to have a high degree of interdependency. In order to sample from this distribution, we use Dynamic Hamiltonian Monte Carlo, a Hamiltonian Monte Carlo (HMC) sampler available in Stan [35]. HMC is a Markov chain Monte Carlo approach that due to possessing energy conserving properties is able to take large steps between individual states while maintaining a high acceptance rates. This makes it efficient at sampling from moderately high dimensional posterior distributions with differentiable likelihoods, while requiring a much lower number of iterations. We implemented the model and inference method in a R package which is available at https://github.com/dhelekal/ResistPhy/. All results shown used 4 chains with 2000 iterations for warmup and 2000 iterations for sampling. For all model parameters and all analysis the bulk effective sample size (bulk-ESS) was always greater than 500, and all *R* statistics were lower than 1.05 [36], values that indicate no issues with mixing. We also checked that there were no divergent transitions at least during the sampling phase.

### Use of simulated and real datasets

For all simulations we use a stochastic, discrete state-space version of the multi-lineage SIS in Equation 1. The system is simulated using tau-leaping [37]. More specifically we consider a scenario with three lineages simulated over the course of 19 years. Two lineages are set to be susceptible and thus unaffected by antibiotic usage fluctuations and one is set to be resistant. The first lineage aims to represent the unobserved bulk of the population and thus is set to start at much higher prevalence. Conditional on the trajectories of the two lineages, we sample phylogenies under Kingman’s coalescent with varying effective population size *N*_e_(*t*) following Equation 6 conditional on the trajectories [28]. The parameters for the simulation were selected as to consistently provide a reasonable range of plausible behaviours so that resistant lineages would reach prevalence with orders of magnitude between 10^2^ to 10^4^.

A total of 1102 genomes were collected between 2000 and 2013 by the CDC Gonococcal Isolate Surveillance (GISP) Project [38]. A maximum-likelihood phylogeny was computed using PhyML [39], which was corrected for recombination using ClonalFrameML [40] and dated using BactDating [24]. This dated phylogeny is the same as previously used in an analysis of hidden population structure [41]. The distribution of primary antimicrobial drugs used to treat gonorrhoea among participants of the GISP project between 1988 and 2019 was obtained from the GISP reports available at https://www.cdc.gov/std/statistics/archive.htm. Note that usage of ciprofloxacin and ofloxacin were combined into a single fluoroquinolone category. All the data and code used in the simulated and real datasets analyses is available at https://github.com/dhelekal/ResistPhy/tree/main/run.

## RESULTS

### Detailed analysis of a single simulated dataset

To validate the performance of this model we first resort to simulation from a 3-lineages stochastic SIS with population size *N* (*t*), transmission rate *β*(*t*) and antimicrobial usage function *u*(*t*) varying over the past 20 years, as illustrated in Figure 1. The first two lineages are susceptible and thus unaffected by fluctuations in antimicrobial usage, whereas the third lineage is resistant and therefore affected. The first lineage represents the bulk of the susceptible lineages and is thus left unobserved. The remaining two lineages represent the observed lineages, susceptible and resistant, respectively. The per-day recovery rate of the sensitive lineage was set to *γ*_*s*_ = 1/60, the fitness cost of resistance to *q*_*U*_ = 1.25 and the fitness benefit of resistance to *q*_*T*_ = −2.7. From each of these two observed lineages, a dated phylogeny with 200 leaves was simulated. The sampling dates were randomly assigned to one of the first six years, with the relative probability of a particular year being chosen proportional to the total prevalence in that year. We performed inference on this simulated dataset; the traces are shown in Figure S1 and the posterior distribution of the kernel parameters in Figure S2. The prevalence and reproduction number *R*(*t*) of both the susceptible and resistant lineages are shown in Figure 2. As expected, the inferred values followed the correct values used in the simulation. The inferred values of the susceptible lineage recovery rate *γ*_*s*_ and the cost and benefit of resistance *q*_*U*_ and *q*_*T*_ were also found to be close to their correct values, as shown in Figure 3. The posterior distribution of *γ*_*s*_ was almost identical to the prior, which was centered on the correct value 1/60, reflecting the fact that the data is uninformative about this parameter and stressing the importance of using an informative prior. There was a strong negative correlation between the inferred values of *q*_*U*_ and *q*_*T*_, as expected since these two parameters play opposite roles in the overall fitness of the resistant lineage relative to the sensitive lineage. Nevertheless, we detected both the cost and the benefit associated with resistance, since the ranges of inferred values for *q*_*U*_ and *q*_*T*_ were respectively above and below one, contrary to their log-normal priors with mean one (Figure 3). Finally, we computed the posterior predictive distribution [42] for the number of ancestral lineages through time *A*(*t*) and compared this with the input phylogenetic data (Figure S3). The data and posterior predictive trajectories were similar, indicating a good fit of the model to the data as indeed would be expected here since the same model was used for simulation and inference.

**Figure 1:**
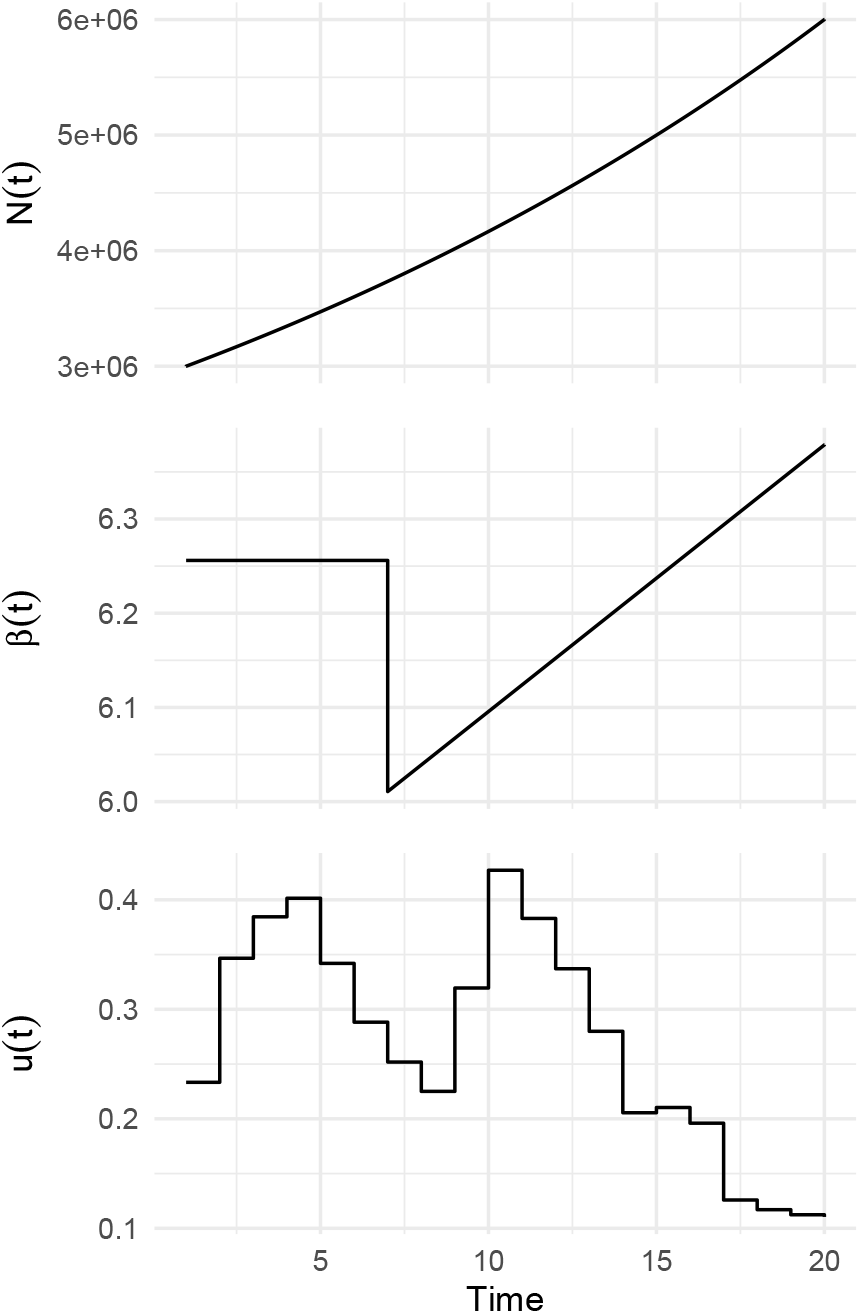
Host population size function *N* (*t*), transmission rate over time *β*(*t*) and antibiotic usage function *u*(*t*) used in the simulated datasets.

**Figure 2:**
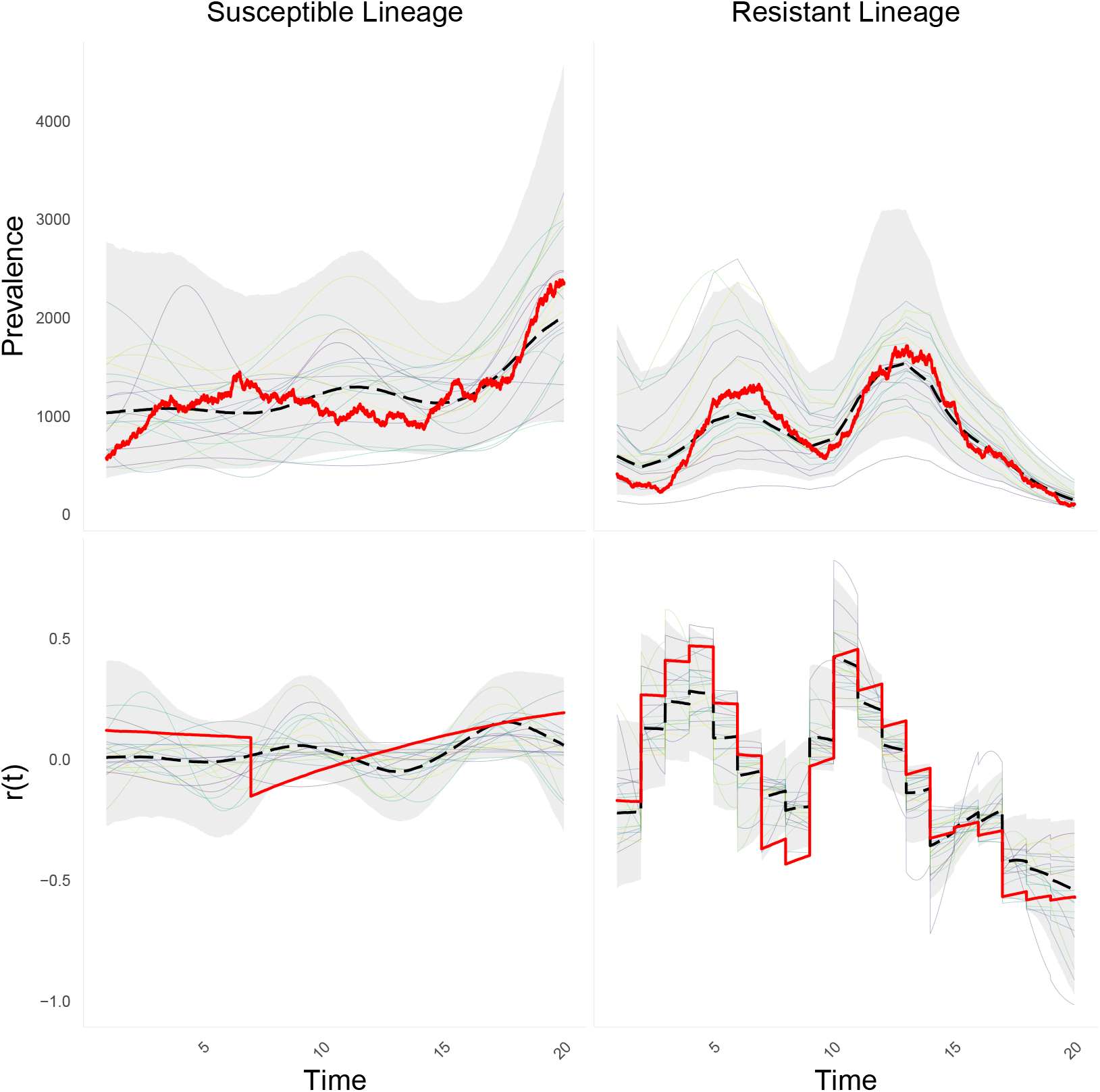
Posterior summary of dynamics for the sensitive (left) and resistant (right) lineages, showing prevalence (top) and reproduction number (bottom). Bold solid red lines indicates simulated values. Posterior median in bold dashed black line. Shaded bands indicate 95% posterior credible intervals. Solid light lines represent posterior draws.

**Figure 3:**
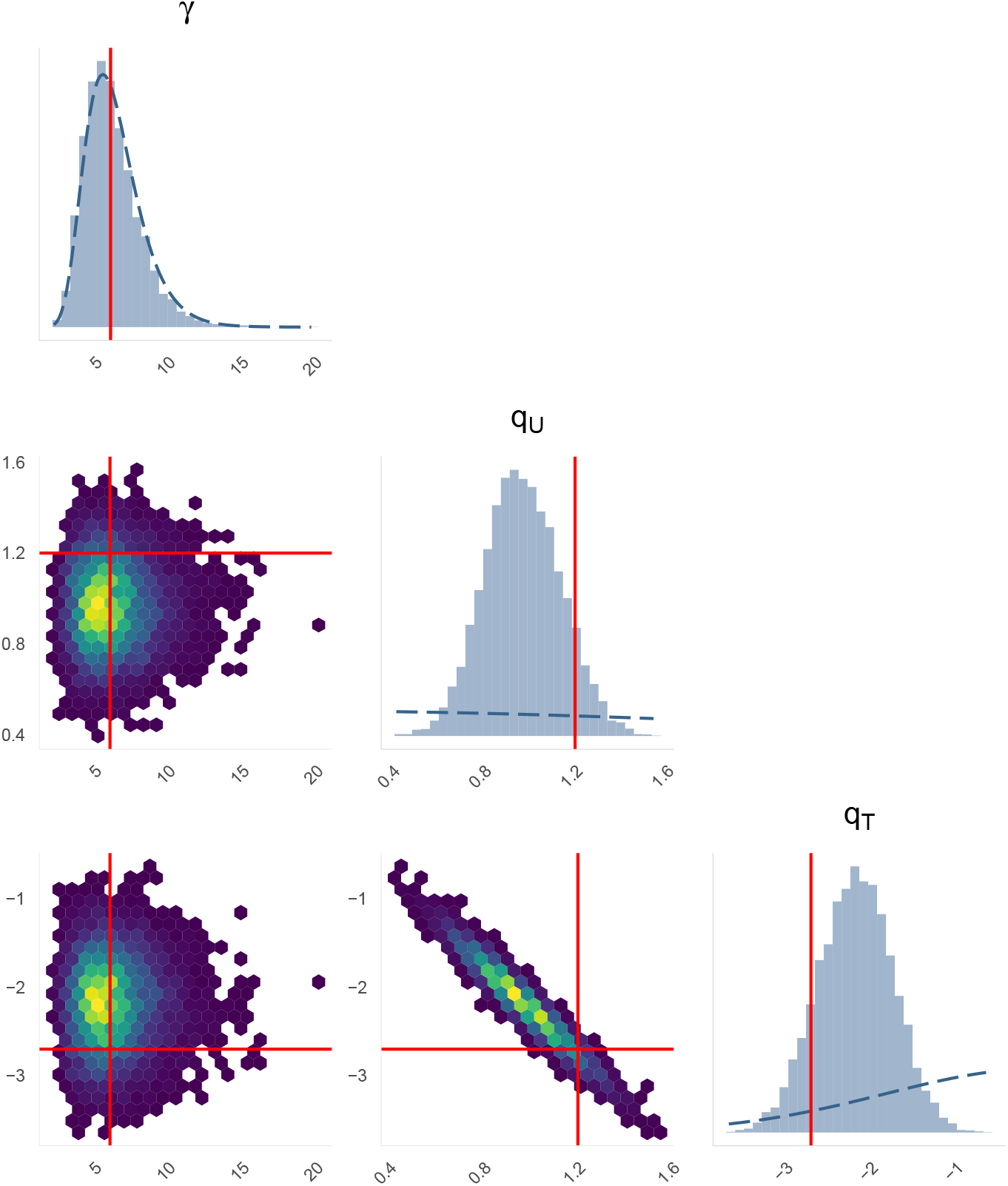
Marginal and joint posterior distributions for the recovery rate of the sensitive lineage (*γ*_*s*_), fitness cost (*q*_*U*_) and fitness benefit (*q*_*T*_) of resistance. Bold red solid lines indicate simulation values. Bold blue dashed lines indicate prior density values.

### Benchmark using multiple simulated datasets

We repeated the same application of our inference method to data simulated in the same conditions as described above and illustrated in Figure 1, except the values of the fitness cost and benefit of resistance were varied. A total of 50 simulated datasets were generated and analysed, with the fitness cost *q*_*U*_ increasing linearly from 1 to 1.2, and the fitness benefit *q*_*T*_ decreasing linearly from 1 to 0.5. The prevalence of the susceptible and resistant lineages in these simulations are shown in Figure S4. The results of inference are illustrated in Figure 4 and show that in almost all cases the posterior 95% credible intervals covered the correct values of the fitness cost and benefit of resistance used in the simulations.

**Figure 4:**
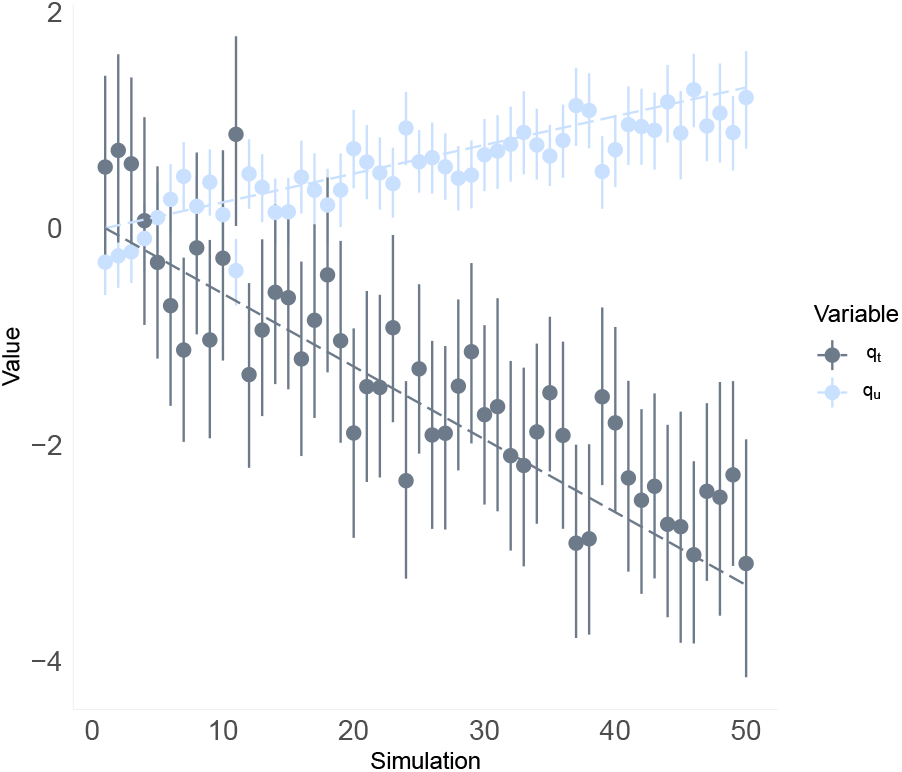
Inferred parameters versus correct values. A total of 50 simulated datasets were generated, with decreasing values of *q*_*T*_ and increasing values of *q*_*U*_ as shown by the dotted lines in grey and blue, respectively. For each simulated dataset we applied our inference method. The grey and blue dots show the mean inferred values of *q*_*T*_ and *q*_*U*_, respectively, with vertical bars representing the 95% credible intervals for both parameters.

### Application to fluoroquinolone resistant *N. gonorrhoeae* in USA

We demonstrate the use of our model and inferential framework by estimating the cost and benefit of fluoroquinolone resistance in *N. gonorrhoeae*. Based on the 1102 genomes collected between 2000 and 2013 by the CDC Gonococcal Isolate Surveillance Project [38], a recombination-corrected tree was constructed using ClonalFrameML [40] and dated using BactDating [24]. As there are two major fluoroquinolone resistant lineages present in this phylogeny [38], we decided to do a comparative study. The two fluoroquinolone resistant lineages and one fluoroquinolone susceptible lineage were selected based on similar resistance profiles against other relevant antibiotics. By inspecting the antibiotic usage data and the resistance profiles for the the three lineages (Figure 5) we can see that the resistance profiles match for antimicrobials that were in use as primary treatment at significant levels after 1995. As such this is the year we set as the analysis start date (*t*_min_ = 1995) and the end date is the date when the last genomes were collected (*t*_max_ = 2013). Note that a subclade within the susceptible lineage that displayed a *de novo* gain of resistance to cefixime has been removed. The prior mean for the per-day recovery rate for the susceptible lineage was set to *γ*^*^ = 1/90 based on previous gonorrhoea modelling studies [10, 43, 44].

**Figure 5:**
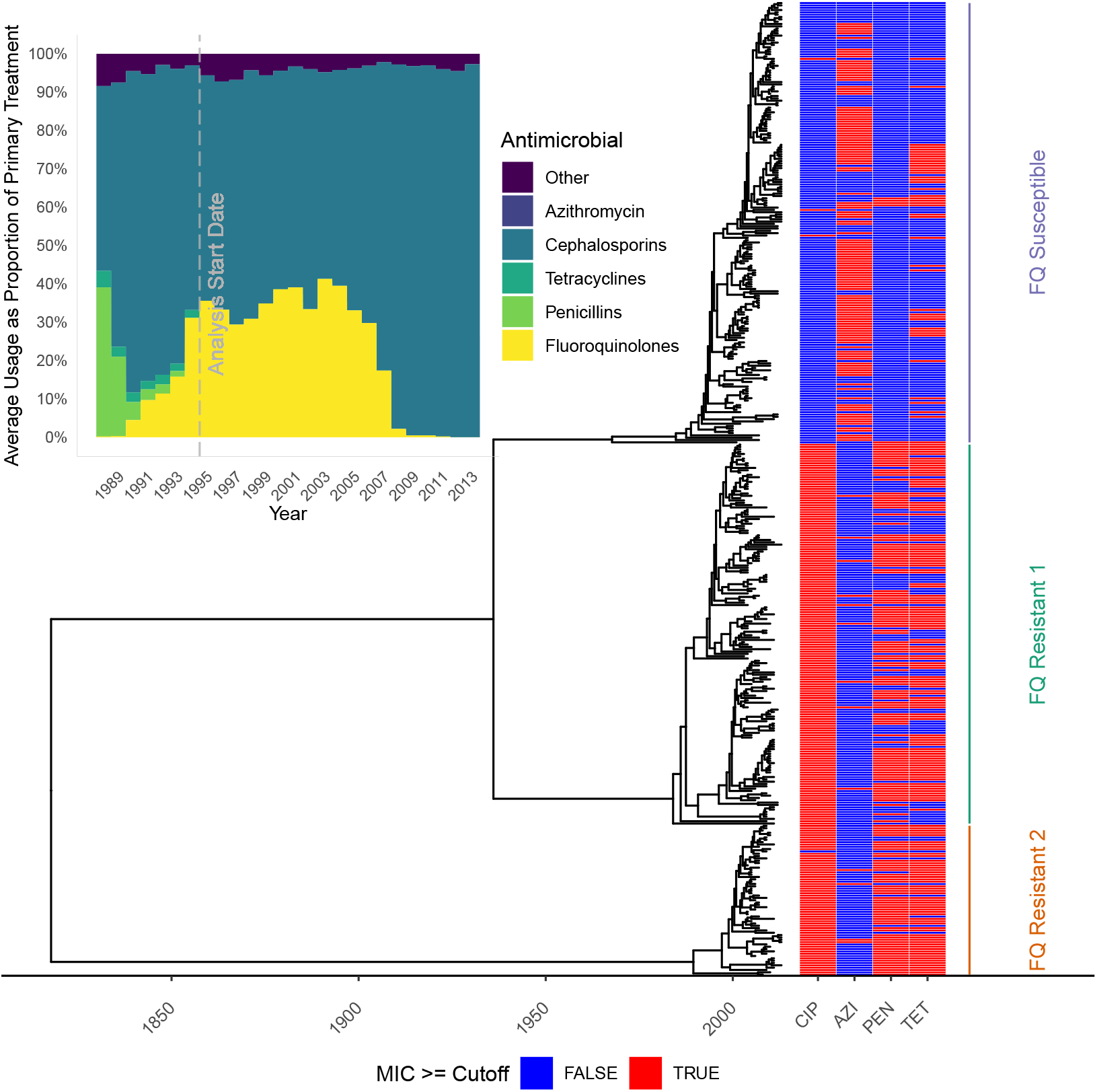
Antibiotic usage data and phylogeny used for the application to fluoroquinolone resistant *N. gonorrhoeae*.

We performed inference for this dataset; the traces are shown in Figure S5 and the posterior distribution of kernel parameters in Figure S6. Figure 6 depicts the summary of posterior latent transmission dynamics for the two resistant lineages, whereas Figure S7 shows the same for the susceptible lineage. The two resistant lineages have similar dynamics, with a peak in prevalence around 2007, which corresponds to the moment when fluoroquinolone use dropped (Figure 5). Figure 7 depicts the marginal and joint posterior distributions for the resistance parameters *q*_*U*_ and *q*_*T*_ for both resistant lineages. This is consistent with there being both a cost and benefit to fluoroquinolone resistance for both lineages, since both *q*_*T*_ and *q*_*U*_ are respectively localised below 1 and above 1, with high posterior probability. It is noteworthy that while both of these lineages come from distinct genetic background, their resistance profile is qualitatively very similar, indicating both of these lineages faced similar selective pressures and neither seems to have successfully adapted to overcome the fitness cost associated with fluoroquinolone resistance. We used a posterior predictive approach to ensure that the model can explain the data appropriately [42]. Posterior predictive trajectories for the function of ancestral lineages through time *A*(*t*) were simulated and found to be very similar to the ones implied by the phylogenetic data (Figure S8).

**Figure 6:**
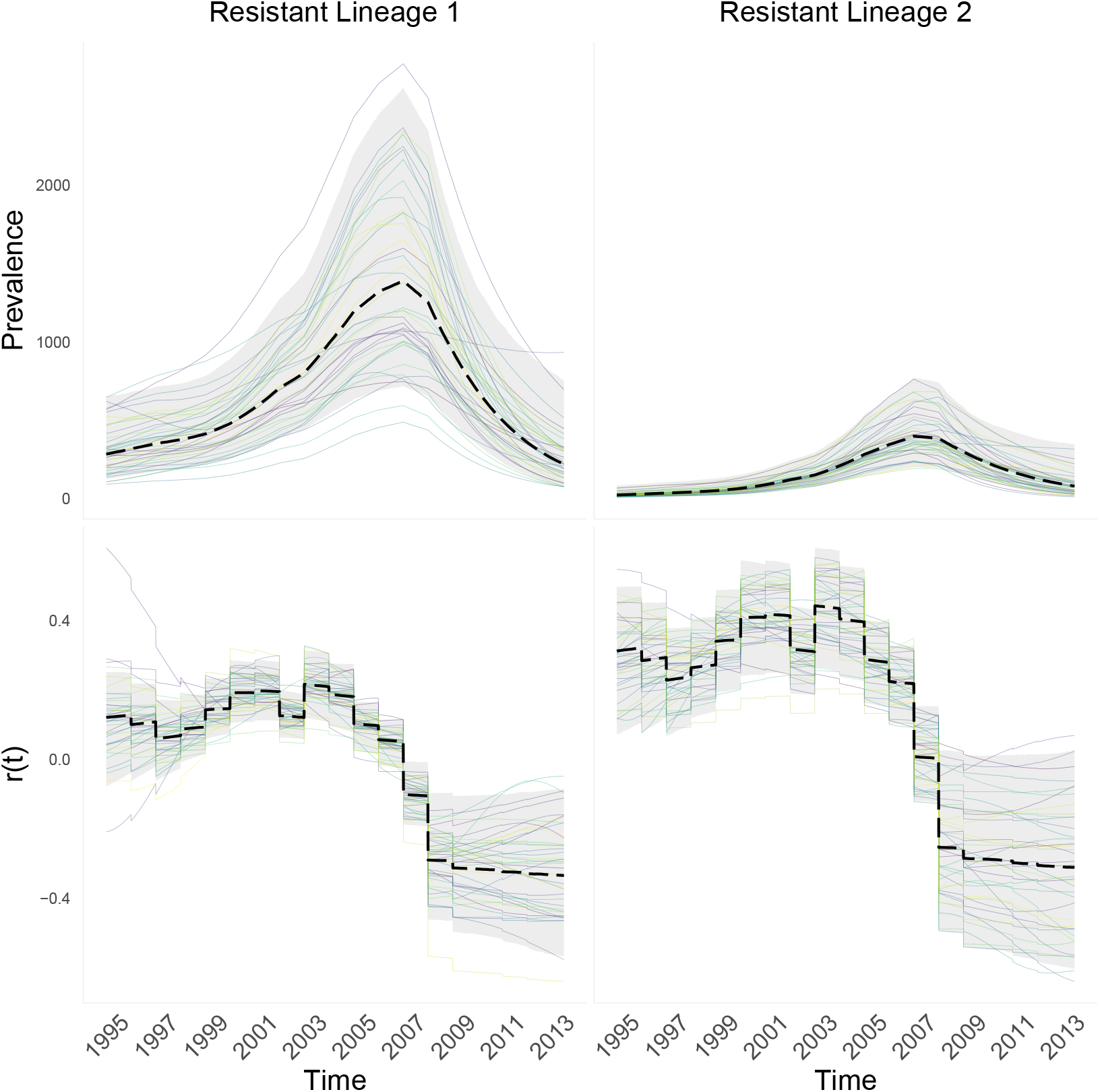
Posterior epidemic dynamics for both fluoroquinolone resistant lineages of *N. gonorrhoeae*.

**Figure 7:**
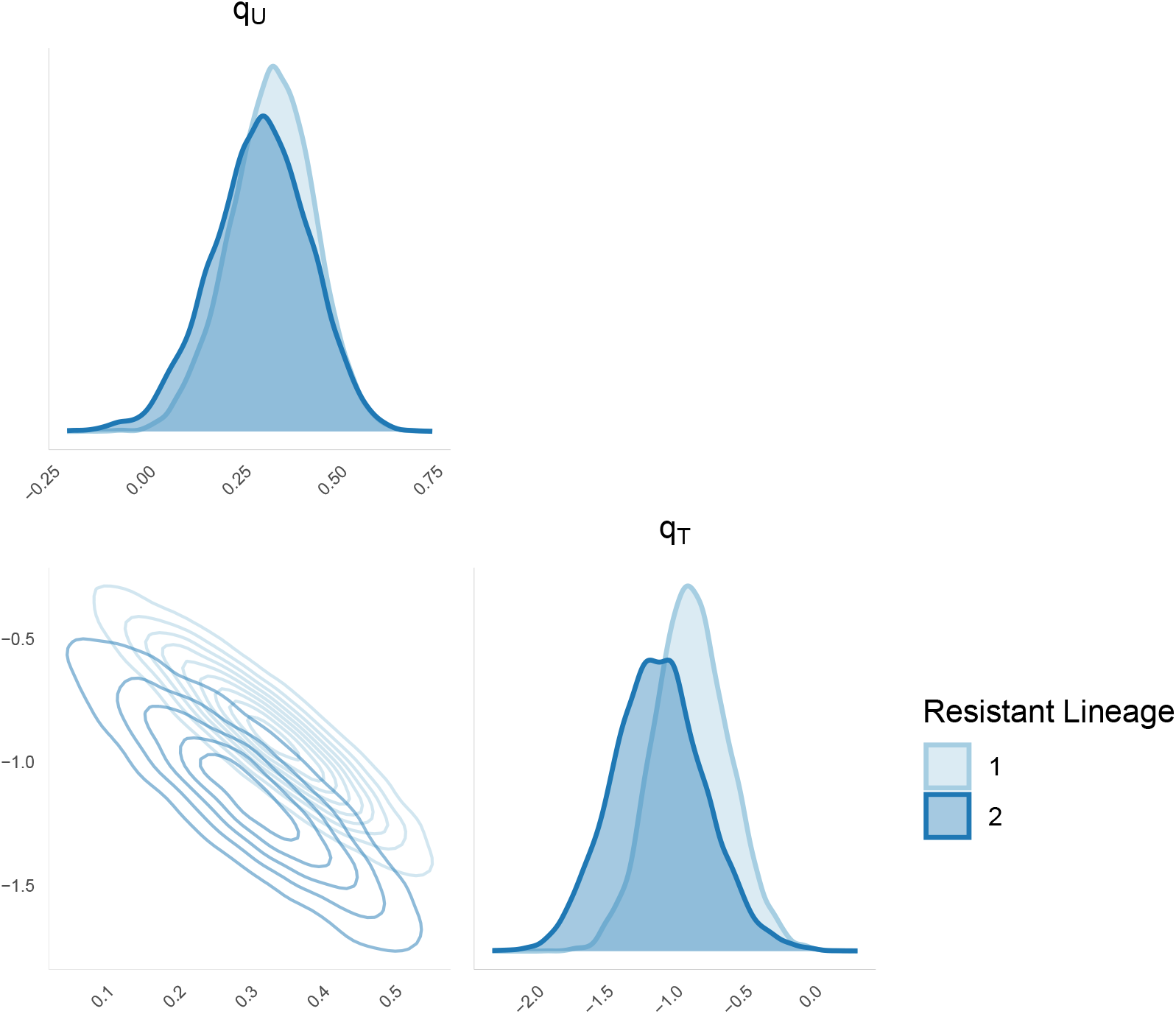
Marginal and joint posterior distribution for the cost (*q*_*U*_) and benefit (*q*_*T*_) of both fluoroquinolone resistant lineages of *N. gonorrhoeae*.

Under the assumption of perfect competition between lineages, if we want to ensure to that a resistant lineage cannot establish, and its proportion decays sufficiently fast, we fix a decay factor *c* > 0 and aim to ensure that the growth rate of the resistant lineage is *c* units lower than that of the sensitive lineage, that is *r*_*s*_(*t*) −*r*_*r*_(*t*) > *c*. Note that *r*(*t*) is the growth rate through time, not *R*(*t*), the time varying reproduction number. We choose to work with growth rates as these are less sensitive to susceptible recovery rate mispecification. Given that the lineages have the same transmission rate function *b*(*t*), this condition is equivalent to *γ*_*s*_(*t*) − *γ*_*r*_(*t*) > *c*, and using the definition of *γ*_*r*_(*t*) from Equation 3, this is equivalent to *u*(*t*)*q*_*T*_ + (1− *u*(*t*))*q*_*U*_ > *c*. We use this to estimate posterior probabilities the differences in growth rates between the susceptible lineage and each of both resistant lineages exceed *c* as shown in Figure 8. In order to be 95% certain that the resistant lineages remain at a lower fitness than the susceptible lineage, fluoroquinolone should not be prescribed to more than ∼20% and ∼15% of infected individuals, for resistant lineages 1 and 2, respectively.

**Figure 8:**
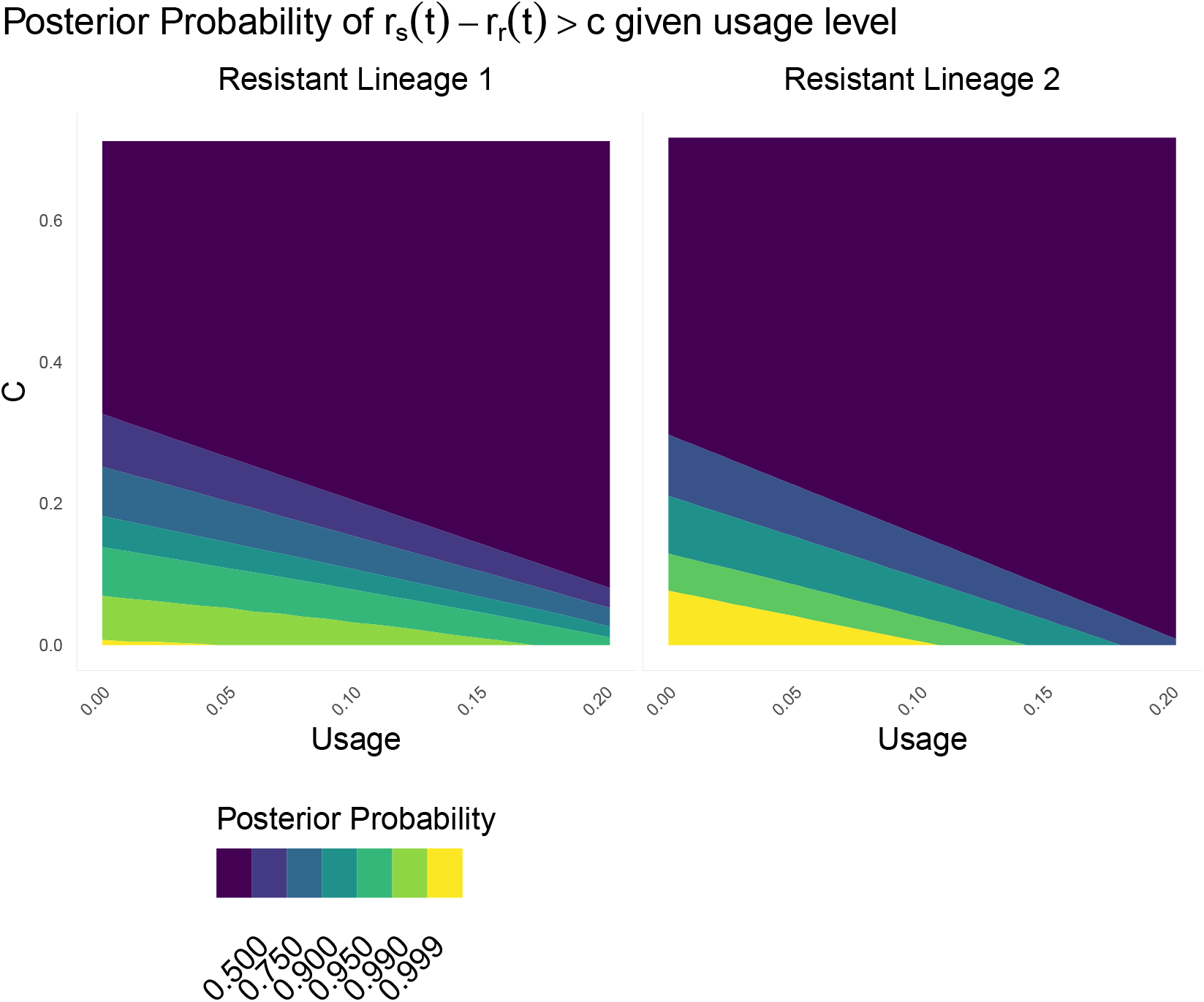
Posterior probabilities of *R*_*r*_(*t*)*/R*_*s*_(*t*) < *C* given usage *u*(*t*) in the x-axis and value of *C* in the y-axis, for both fluoroquinolone resistant lineages of *N. gonorrhoeae*.

## DISCUSSION

A bacterial pathogen lineage that is resistant to a given antibiotic incurs both a fitness cost and a fitness benefit compared to similar susceptible lineages [8]. When the antibiotic is used extensively, the benefit is likely to be greater than the cost. In that case, a resistant lineage has a selective advantage over susceptible lineages, and therefore grows at a faster rate. Conversely, if the antibiotic is used rarely or not at all, the benefit is likely to become smaller than the cost, which will lead to the resistant lineage decreasing in frequency. Estimating these parameters is therefore of primary importance to determine how antibiotics should be prescribed without causing an increase in resistance [9]. Here, we have shown how genome sequencing data coupled with data on antibiotic prescriptions can be used for this purpose, following on previous work that demonstrated the link between epidemic dynamics and phylogenetics [13, 19, 20, 28]. By comparing the phylodynamic trajectories of susceptible and resistant lineages, and relating them with a known function of antibiotic use, we show that it is possible to estimate separately the parameters corresponding to the fitness cost and benefit of resistance. In particular, we reanalysed a large published collection of *N. gonorrhoeae* genomes [38]. We were able to infer these parameters for two lineages of *N. gonorrhoeae* resistant to fluoroquinolones, and found similar estimates of cost and benefit in both (Figure 7). We were able to use this knowledge to make recommendations on antibiotic stewardship of fluoroquinolones (Figure 8).

Dated phylogenies for both susceptible and resistant lineages are needed as input into our method. Several software tools can be used to produce this either from a sequence alignment, for example BEAST [22] and BEAST2 [23], or from an undated phylogeny, for example treedater [45] and BactDating [24]. Building such a dated phylogeny requires either the population to be measurably evolving over the sampling period [46, 47], or a previous estimate of the molecular clock rate [48]. Another input required by our method is the antibiotic usage function over a relevant timeframe and geographical location. This may not always be available in all historical contexts, but efforts are increasingly being made to capture this data [49]. Finally, our method requires an informative prior of the recovery rate for the susceptible lineage (see Table 1), since this is typically not identifiable from the data, as in many similar compartmental epidemic models [50]. This prior needs to be chosen carefully depending on the infectious disease under study and based on the existing scientific literature.

Our inferential methodology is based on a well-defined and relatively simple epidemic model (Equation 4) which means making a number of assumptions the validity of which was considered before performing our analysis. Our model assumes multiple-lineage pathogen dynamics driven by person-to-person transmission in a well mixed host population in the absence of any significant population structure, so that there is perfect competition between lineages. It also assumes that individuals become infectious as soon as they are infected, that their infectiousness remains constant until they recover, after which they become susceptible again without any immunity being gained. This list of relatively strong assumptions may seem to preclude application to any real infectious disease, but they are necessary to obtain a model under which inference can be performed. Furthermore, violation of some of these assumptions does not necessarily invalidate the results of inference. For example, if infection causes immunity, this will effectively reduce the number *S*(*t*) of susceptible individuals (Equation 2), but this number is not assumed to be constant in our model. In fact both the size *N* (*t*) of the host population and the number *S*(*t*) of susceptible individuals are integrated out as part of our parameterisation in terms of the function *b*(*t*) (cf Equation 4), so the inference is robust as long as the immunity conferred applies to all lineages under study. Likewise the assumption of an unstructured population may seem problematic, including in our application to *N. gonorrhoeae* throughout the USA, but for anything other than small local outbreaks the genomes available for analysis are sparsely sampled from the whole infected population [51]. In these conditions, any effect of the host population structure on phylodynamics is likely to be insignificant as long as an effective rather than actual number of infections is considered [52, 53].

The compatibility of our model with the phylogenetic data under analysis can be tested using posterior predictive distribution checks (Figures S3 and S8). If these tests fail, or if the model assumptions are thought to be inappropriate, a solution may be to resort to other methods that postprocess a dated phylogeny [25] but make less assumptions, at the cost of not inferring directly the parameters of resistance. Alternative approaches includes non-parametric methods that detect differences in the branching patterns in different lineages [41, 54] as well as methods parameterised in terms of the pathogen population size growth rather than underlying epidemiological drivers [15, 55]. However, our model-based approach is both general and flexible, so that we expect it to be applicable in many settings using our software implementation which is available at https://github.com/dhelekal/ResistPhy/. We believe that this methodology, applied to the increasingly large genomic databases on many bacterial pathogens, will help quantify the exact link between antibiotic usage and resistance and therefore provide a much-needed evidence basis for the design of future antibiotic prescription strategies [9, 56, 57].

## ACKNOWLEDGMENTS

We acknowledge funding from the National Institute for Health Research (NIHR) Health Protection Research Unit in Genomics and Enabling Data. This work was supported by the UK Engineering and Physical Sciences Research Council (EPSRC) grant EP/S022244/1 for the EPSRC Centre for Doctoral Training in Mathematics for Real-World Systems II.

## Notes

https://github.com/dhelekal/ResistPhy/

### Competing Interest Statement

The authors have declared no competing interest.

### Summary of Updates

Clarification of Methods sections, better contextualisation in Introduction and Discussion.

## References

[1] CDC, 2013 Antibiotic resistance threats in the United States, 2013. Current p. 114. ISSN 10985530. (doi:CS239559-B).

[2] O’Neill, J., 2016 Tackling drug-resistant infections globally: final report and recommendations. London: Wellcome Trust\&HM Government.

[3] Murray, C. J., Ikuta, K. S., Sharara, F., Swetschinski, L., Aguilar, G. R., Gray, A., Han, C., Bisignano, C., Rao, P., Wool, E. et al., 2022 Global burden of bacterial antimicrobial resistance in 2019: a systematic analysis. The Lancet 399, 629–655.

[4] Clatworthy, A. E., Pierson, E. & Hung, D. T., 2007 Targeting virulence: a new paradigm for antimicrobial therapy. Nat. Chem. Biol. 3, 541–548. ISSN 1552-4450. (doi:10.1038/nchembio.2007.24).

[5] Bonhoeffer, S., Lipsitch, M. & Levin, B. R., 1997 Evaluating treatment protocols to prevent antibiotic resistance. Proc Natl Acad Sci U S A 94, 12106–12111. ISSN 0027-8424 (Print). (doi:10.1073/pnas.94.22.12106).

[6] Austin, D. J., Kristinsson, K. G. & Anderson, R. M., 1999 The relationship between the volume of antimicrobial consumption in human communities and the frequency of resistance. PNAS 96, 1152–1156. ISSN 0027-8424. (doi:10.1073/pnas.96.3.1152).

[7] Spicknall, I. H., Foxman, B., Marrs, C. F. & Eisenberg, J. N. S., 2013 A modeling framework for the evolution and spread of antibiotic resistance: Literature review and model categorization. Am. J. Epidemiol. 178, 508–520. ISSN 00029262. (doi:10.1093/aje/kwt017).

[8] Andersson, D. I. & Levin, B. R., 1999 The biological cost of antibiotic resistance. Curr. Opin. Microbiol. 2, 489–493. ISSN 13695274. (doi:10.1016/S1369-5274(99)00005-3).

[9] Andersson, D. I. & Hughes, D., 2010 Antibiotic resistance and its cost: is it possible to reverse resistance? Nat. Rev. Microbiol. 8, 260–271. ISSN 1740-1526. (doi:10.1038/nrmicro2319).

[10] Whittles, L. K., White, P. J. & Didelot, X., 2017 Estimating the fitness benefit and cost of cefixime resistance in Neisseria gonorrhoeae to inform prescription policy: A modelling study. PLoS Med. 14, e1002416. (doi:10.1371/journal.pmed.1002416).

[11] Rubin, D. H., Ma, K. C., Westervelt, K. A., Hullahalli, K., Waldor, M. K. & Grad, Y. H., 2022 Variation in supplemental carbon dioxide requirements defines lineage-specific antibiotic resistance acquisition in neisseria gonorrhoeae. bioRxiv (doi:10.1101/2022.02.24.481660). Publisher: Cold Spring Harbor Laboratory eprint: https://www.biorxiv.org/content/early/2022/02/25/2022.02.24.481660.full.pdf.

[12] Lehtinen, S., Blanquart, F., Lipsitch, M., Fraser, C. & Collaboration, w. t. M. P., 2019 On the evolutionary ecology of multidrug resistance in bacteria. PLOS Pathogens 15, e1007763. ISSN 1553-7374. (doi:10.1371/journal.ppat.1007763). Publisher: Public Library of Science.

[13] Pybus, O. G. & Rambaut, A., 2009 Evolutionary analysis of the dynamics of viral infectious disease. Nat. Rev. Genet. 10, 540–50. ISSN 1471-0064. (doi:10.1038/nrg2583).

[14] Volz, E. M. & Frost, S. D. W., 2014 Sampling through time and phylodynamic inference with coalescent and birth – death models. J. R. Soc. Interface 11, 20140945.

[15] Volz, E. M. & Didelot, X., 2018 Modeling the Growth and Decline of Pathogen Effective Population Size Provides Insight into Epidemic Dynamics and Drivers of Antimicrobial Resistance. Syst. Biol. 67, 719–728. ISSN 1063-5157. (doi:10.1093/sysbio/syy007).

[16] Kuühnert, D., Kouyos, R., Shirreff, G., Pe?cerska, J., Scherrer, A. U., Büoni, J., Yerly, S., Klimkait, T., Aubert, V., Guünthard, H. F. et al., 2018 Quantifying the fitness cost of HIV-1 drug resistance mutations through phylodynamics. PLOS Pathogens 14, e1006895. ISSN 1553-7374. (doi: 10.1371/journal.ppat.1006895). Publisher: Public Library of Science.

[17] Pecerska, J., Kuühnert, D., Meehan, C. J., Coscolla, M., de Jong, B. C., Gagneux, S. & Stadler, T., 2021 Quantifying transmission fitness costs of multi-drug resistant tuberculosis. Epidemics 36, 100471. ISSN 1755-4365. (doi:10.1016/j.epidem.2021.100471).

[18] Ho, S. Y. W. & Shapiro, B., 2011 Skyline-plot methods for estimating demographic history from nucleotide sequences. Molecular Ecology Resources 11, 423–434. ISSN 1755098X. (doi: 10.1111/j.1755-0998.2011.02988.x).

[19] Volz, E. M., Kosakovsky Pond, S. L., Ward, M. J., Leigh Brown, A. J. & Frost, S. D. W., 2009 Phylodynamics of infectious disease epidemics. Genetics 183, 1421–30. ISSN 1943-2631. (doi:10.1534/genetics.109.106021).

[20] Dearlove, B. & Wilson, D., 2013 Coalescent inference for infectious disease: Meta-analysis of hepatitis C. Philosophical Transactions of the Royal Society B 368, 20120314.

[21] Didelot, X., Bowden, R., Wilson, D. J., Peto, T. E. A. & Crook, D. W., 2012 Transforming clinical microbiology with bacterial genome sequencing. Nature Reviews Genetics 13, 601–612. ISSN 1471-0056. (doi:10.1038/nrg3226).

[22] Suchard, M. A., Lemey, P., Baele, G., Ayres, D. L., Drummond, A. J. & Rambaut, A., 2018 Bayesian phylogenetic and phylodynamic data integration using BEAST 1.10. Virus Evol. 4, vey016. ISSN 2057-1577. (doi:10.1093/ve/vey016).

[23] Bouckaert, R., Vaughan, T. G., Fourment, M., Gavryushkina, A., Heled, J., Denise, K., Maio, N. D., Matschiner, M., Ogilvie, H., Plessis, L. et al., 2019 BEAST 2.5 : An Advanced Software Platform for Bayesian Evolutionary Analysis. PLoS Comput. Biol. 15, e1006650.

[24] Didelot, X., Croucher, N. J., Bentley, S. D., Harris, S. R. & Wilson, D. J., 2018 Bayesian inference of ancestral dates on bacterial phylogenetic trees. Nucleic Acids Res. 46, e134–e134. ISSN 0305-1048. (doi:10.1093/nar/gky783).

[25] Didelot, X. & Parkhill, J., 2022 A scalable analytical approach from bacterial genomes to epidemiology. Philosophical Transactions of the Royal Society B: Biological Sciences 377, 20210246. (doi:10.1098/rstb.2021.0246).

[26] Allen, L. J., Kirupaharan, N. & Wilson, S. M., 2004 SIS epidemic models with multiple pathogen strains. Journal of Difference Equations and Applications 10, 53–75.

[27] Keeling, M. J. & Rohani, P., 2008 Modeling Infectious Diseases in Humans and Animals. Princeton University Press. ISBN 9780691116174.

[28] Volz, E. M., 2012 Complex population dynamics and the coalescent under neutrality. Genetics 190, 187–201. ISSN 1943-2631. (doi:10.1534/genetics.111.134627).

[29] Griffiths, R. & Tavare, S., 1994 Sampling theory for neutral alleles in a varying environment. Philos. Trans. R. Soc. B 344, 403–410.

[30] Gill, M. S., Lemey, P., Faria, N. R., Rambaut, A., Shapiro, B. & Suchard, M. A., 2013 Improving bayesian population dynamics inference: A coalescent-based model for multiple loci. Mol. Biol. Evol. 30, 713–724. ISSN 07374038. (doi:10.1093/molbev/mss265).

[31] Griffiths, R. C., 2003 The frequency spectrum of a mutation, and its age, in a general diffusion model. Theoretical Population Biology 64, 241–251. ISSN 0040-5809. (doi:10.1016/S0040-5809(03)00075-3).

[32] Etheridge, A., Pfaffelhuber, P. & Wakolbinger, A., 2006 An approximate sampling formula under genetic hitchhiking. The Annals of Applied Probability 16, 685–729. ISSN 1050-5164, 2168-8737. (doi:10.1214/105051606000000114). Publisher: Institute of Mathematical Statistics.

[33] Rasmussen, C. E., 2004 Gaussian Processes in Machine Learning, pp. 63–71. Berlin, Heidelberg: Springer Berlin Heidelberg. ISBN 978-3-540-28650-9. (doi:10.1007/978-3-540-28650-94).

[34] Riutort-Mayol, G., Buürkner, P.-C., Andersen, M. R., Solin, A. & Vehtari, A., 2022 Practical hilbert space approximate bayesian gaussian processes for probabilistic programming. arXiv (doi:10.48550/arXiv.2004.11408).

[35] Carpenter, B., Gelman, A., Hoffman, M. D., Lee, D., Goodrich, B., Betancourt, M., Brubaker, M. A., Guo, J., Li, P. & Riddell, A., 2017 Stan: A probabilistic programming language. J. Stat. Softw. s76. ISSN 15487660. (doi:10.18637/jss.v076.i01).

[36] Vehtarh, A., Gelman, A., Simpson, D., Carpenter, B. & Burkner, P. C., 2021 Rank-Normalization, Folding, and Localization: An Improved R hat for Assessing Convergence of MCMC. Bayesian Anal. 16, 667–718. ISSN 19316690. (doi:10.1214/20-BA1221).

[37] Gillespie, D. T., 2001 Approximate accelerated stochastic simulation of chemically reacting systems. The Journal of Chemical Physics 115, 1716–1733. ISSN 0021-9606. (doi:10.1063/1.1378322). Publisher: American Institute of Physics.

[38] Grad, Y. H., Harris, S. R., Kirkcaldy, R. D., Green, A. G., Marks, D. S., Bentley, S. D., Trees, D., Lipsitch, M., Diseases, I., Health, P. et al., 2016 Genomic epidemiology of gonococcal resistance to extended spectrum cephalosporins, macrolides, and fluoroquinolones in the US, 2000-2013. J. Infect. Dis. 214, 1579–1587. ISSN 0022-1899. (doi:10.1093/infdis/jiw420).

[39] Guindon, S., Dufayard, J.-F., Lefort, V., Anisimova, M., Hordijk, W. & Gascuel, O., 2010 New algorithms and methods to estimate maximum-likelihood phylogenies: Assessing the performance of PhyML 3.0. Systematic biology 59, 307–21. ISSN 1076-836X. (doi:10.1093/sysbio/syq010).

[40] Didelot, X. & Wilson, D. J., 2015 ClonalFrameML: Efficient Inference of Recombination in Whole Bacterial Genomes. PLoS Comput. Biol. 11, e1004041. ISSN 1553-7358. (doi:10.1371/journal.pcbi.1004041).

[41] Volz, E. M., Wiuf, C., Grad, Y. H., Frost, S. D. W., Dennis, A. M. & Didelot, X., 2020 Identification of hidden population structure in time-scaled phylogenies. Systematic Biology 69, 884–896. (doi:10.1093/sysbio/syaa009).

[42] Gelman, A., Meng, X. & Stern, H., 1996 Posterior predictive assessment of model fitness via realized discrepancies. Stat Sinica 6, 733–807.

[43] Fingerhuth, S. M., Bonhoeffer, S., Low, N. & Althaus, C. L., 2016 Antibiotic-Resistant Neisseria gonorrhoeae Spread Faster with More Treatment, Not More Sexual Partners. PLOS Pathog. 12, e1005611. ISSN 1553-7374. (doi:10.1371/journal.ppat.1005611).

[44] Whittles, L. K., White, P. J. & Didelot, X., 2019 A dynamic power-law sexual network model of gonorrhoea outbreaks. PLoS Comput. Biol. 15, e1006748. ISSN 1553-7358. (doi:10.1371/journal.pcbi.1006748).

[45] Volz, E. M. & Frost, S. D. W., 2017 Scalable relaxed clock phylogenetic dating. Virus Evolution 3, vex025.

[46] Drummond, A. J., Pybus, O. G., Rambaut, A., Forsberg, R. & Rodrigo, A. G., 2003 Measurably evolving populations. Trends in Ecology and Evolution 18, 481–488. ISSN 01695347. (doi:10.1016/S0169-5347(03)00216-7).

[47] Biek, R., Pybus, O. G., Lloyd-Smith, J. O. & Didelot, X., 2015 Measurably evolving pathogens in the genomic era. Trends in Ecology & Evolution 30, 306–313. ISSN 0169-5347. (doi:10.1016/j.tree.2015.03.009).

[48] Duchene, S., Holt, K. E., Weill, F.-X., Le Hello, S., Hawkey, J., Edwards, D. J., Fourment, M. & Holmes, E. C., 2016 Genome-scale rates of evolutionary change in bacteria. Microbial Genomics 2, e000094. ISSN 2057-5858. (doi:10.1101/069492).

[49] Curtis, H. J. & Goldacre, B., 2018 Openprescribing: normalised data and software tool to research trends in english nhs primary care prescribing 1998–2016. BMJ open 8, e019921.

[50] Roosa, K. & Chowell, G., 2019 Assessing parameter identifiability in compartmental dynamic models using a computational approach: application to infectious disease transmission models. Theoretical Biology and Medical Modelling 16, 1–15.

[51] Klinkenberg, D., Colijn, C. & Didelot, X., 2019 Methods for Outbreaks Using Genomic Data. In Handbook of Infectious Disease Data Analysis, pp. 245–263. CRC Press.

[52] Nordborg, M., 1997 Structured coalescent processes on different time scales. Genetics 146, 1501–1514. ISSN 0016-6731.

[53] Frost, S. D. W. & Volz, E. M., 2010 Viral phylodynamics and the search for an ‘effective number of infections’. Philosophical Transactions of the Royal Society B 365, 1879–1890. ISSN 0962-8436. (doi:10.1098/rstb.2010.0060).

[54] Dearlove, B. L., Xiang, F. & Frost, S. D. W., 2017 Biased phylodynamic inferences from analysing clusters of viral sequences. Virus Evolution 3, 1–10. (doi:10.1093/ve/vex020).

[55] Helekal, D., Ledda, A., Volz, E., Wyllie, D. & Didelot, X., 2021 Bayesian inference of clonal expansions in a dated phylogeny. Systematic Biology p. syab095. ISSN 1063-5157. (doi:10.1093/sysbio/syab095).

[56] Michael, C. A., Dominey-Howes, D. & Labbate, M., 2014 The Antimicrobial Resistance Crisis: Causes, Consequences, and Management. Frontiers in Public Health 2. ISSN 2296-2565. (doi:10.3389/fpubh.2014.00145).

[57] Holmes, A. H., Moore, L. S., Sundsfjord, A., Steinbakk, M., Regmi, S., Karkey, A., Guerin, P. J. & Piddock, L. J., 2016 Understanding the mechanisms and drivers of antimicrobial resistance. Lancet 387, 176–187. ISSN 1474547X. (doi:10.1016/S0140-6736(15)00473-0).

