## Supplementary Material for "Estimating the fitness cost and benefit of antimicrobial resistance from pathogen genomic data"

#### Supplementary information for detailed analysis of a single simulated dataset

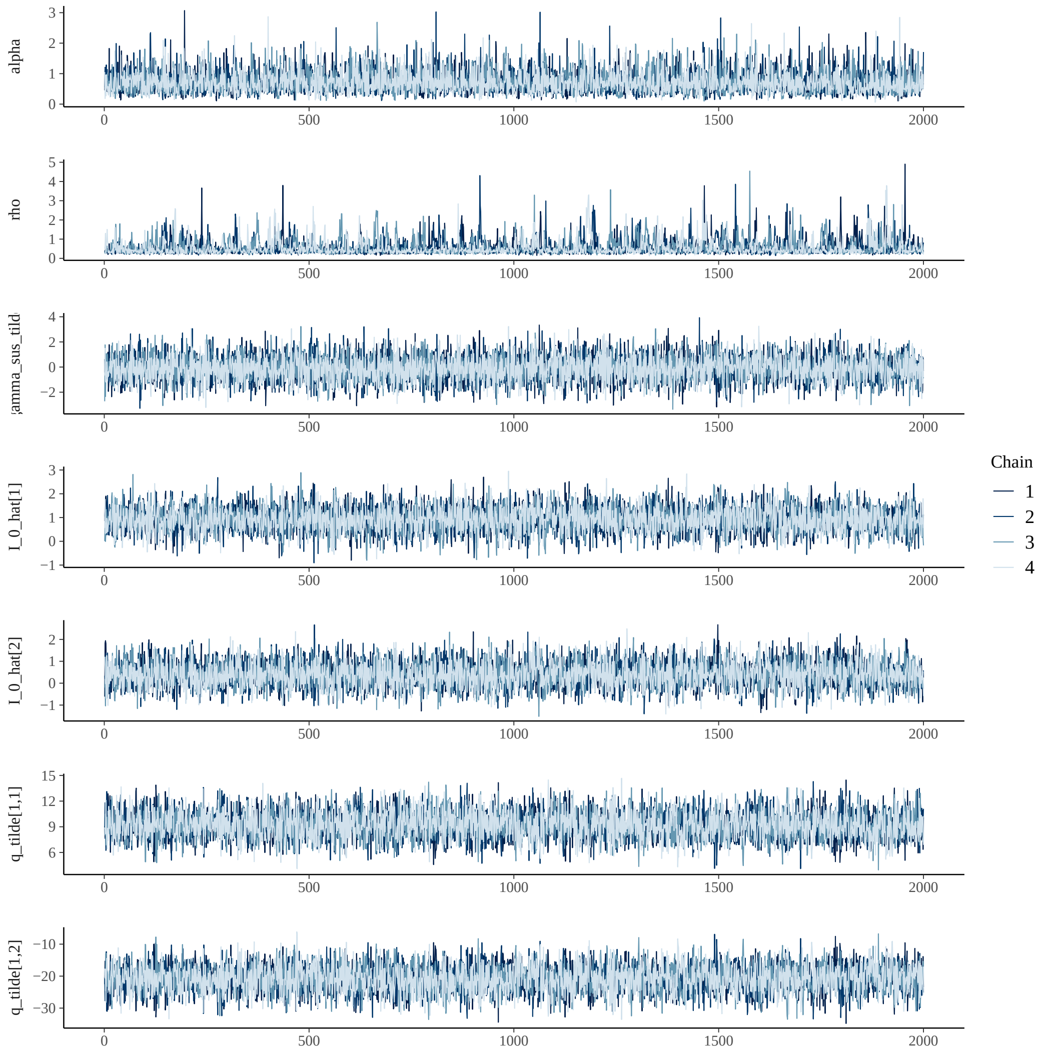


Figure S1: Traces for all parameters excluding GP inducing variables $f_1 : m$ for analysis of single simulated dataset.

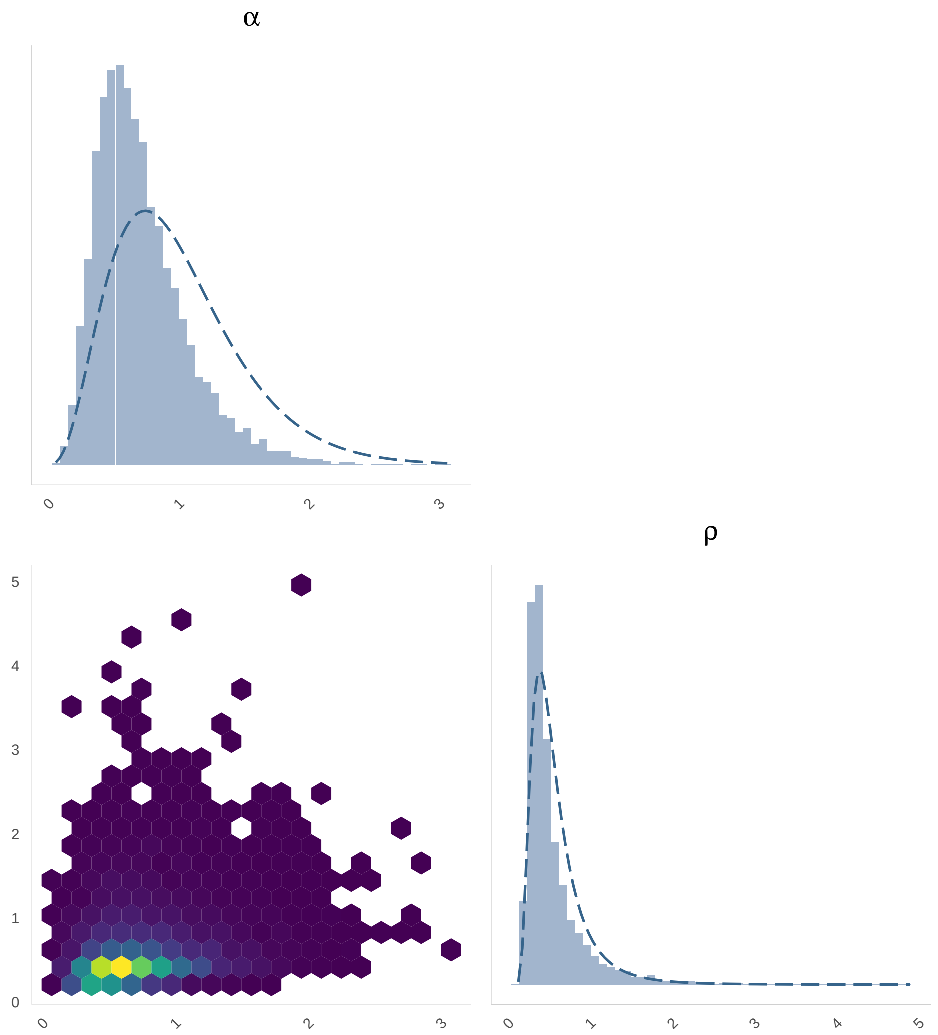


Figure S2: Marginal and joint distributions for hyperparameters $\rho$ and $\alpha$ for the analysis of a single simulated dataset.

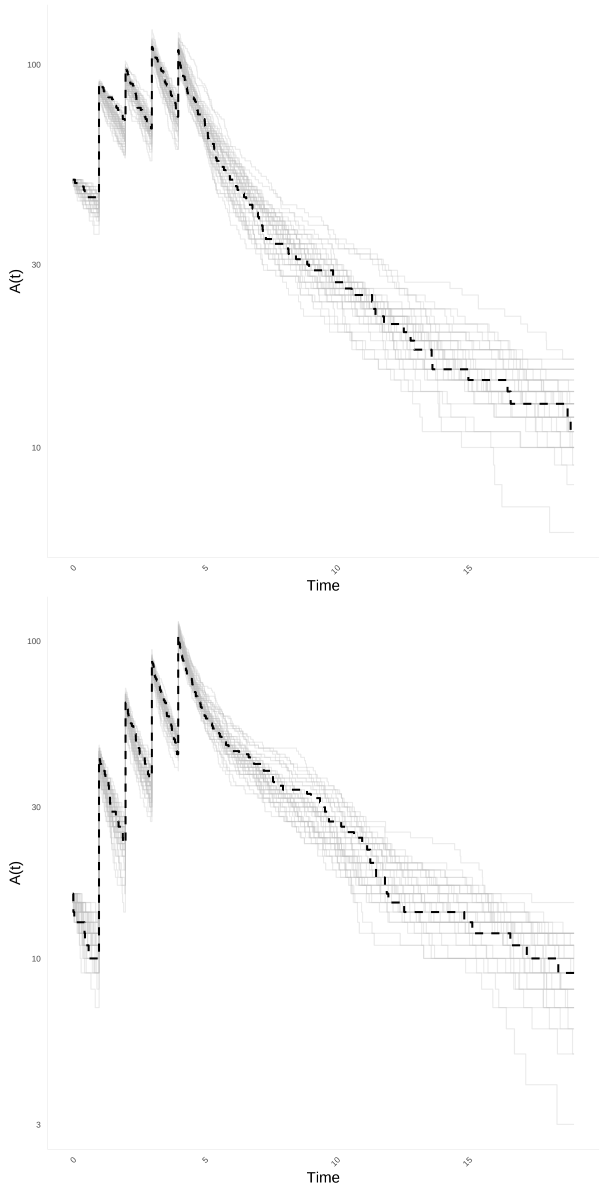


Figure S3: Posterior predictive trajectories of $A(t)$ for the susceptible (top) and resistant (bottom) lineages in the analysis of single simulated dataset.

### Supplementary information for resistance parameter recovery

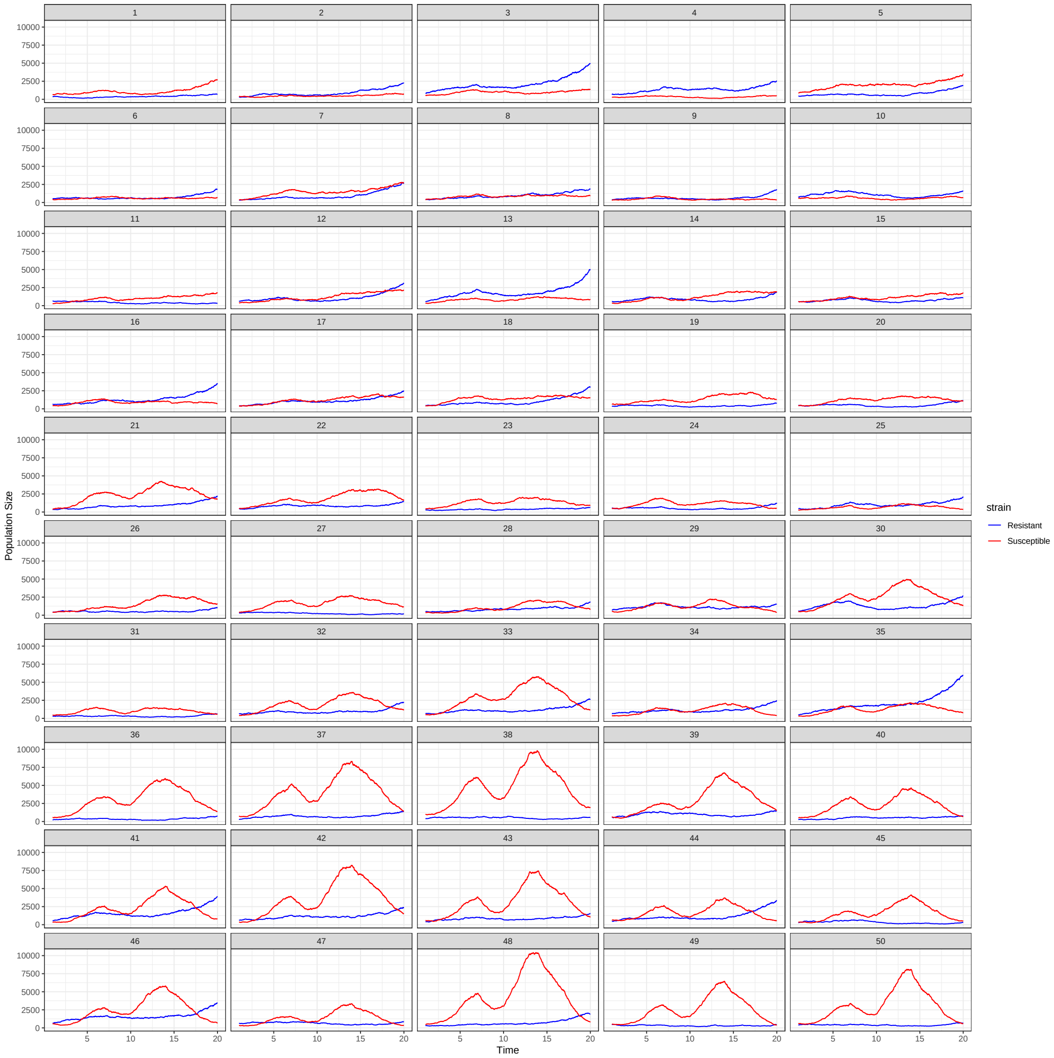


Figure S4: Trajectories of observed lineages used for parameter recovery validation.

### Supplementary information for analysis of *Neisseria gonorrhoeae* dataset

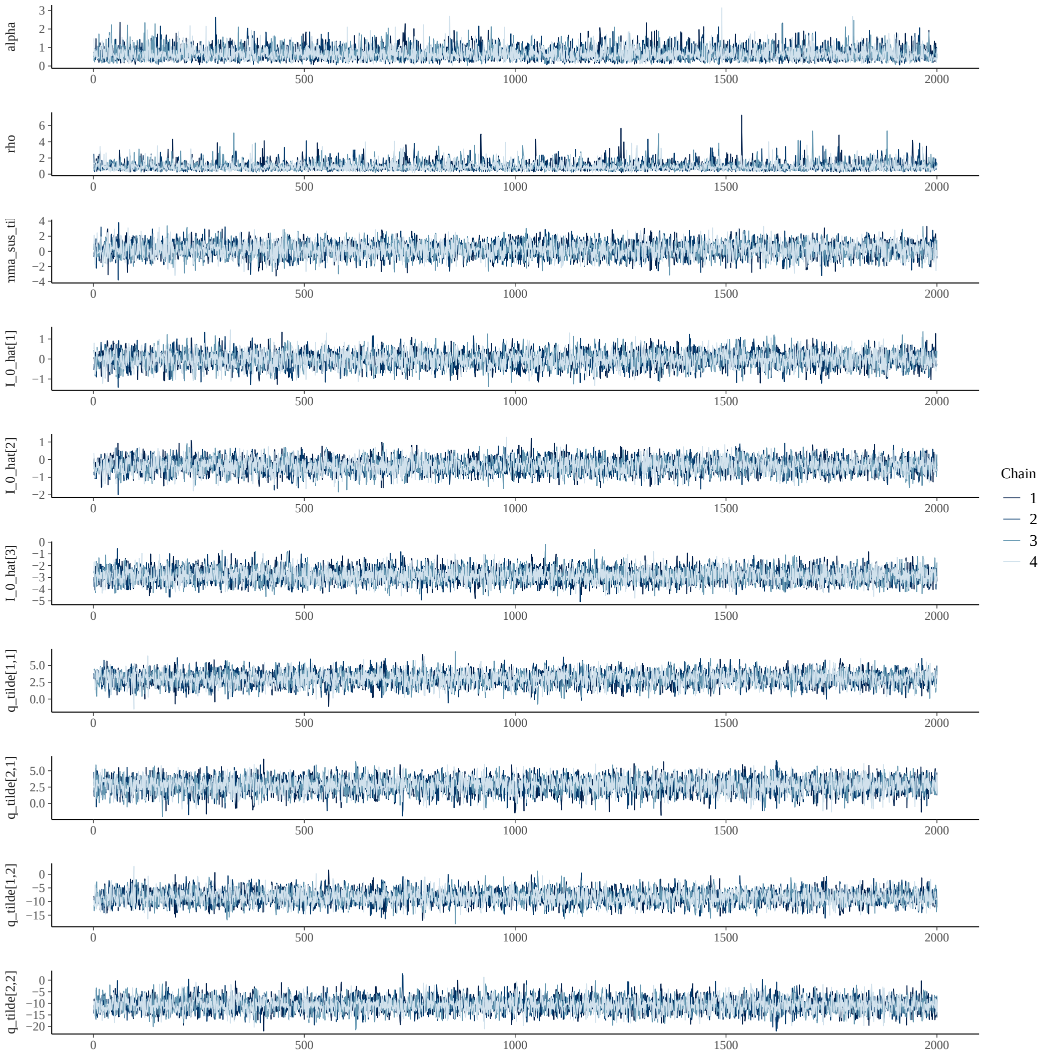


Figure S5: Traces for all parameters excluding GP inducing variables $f_1 : m$ for the analysis of *N. gonorrhoeae*.

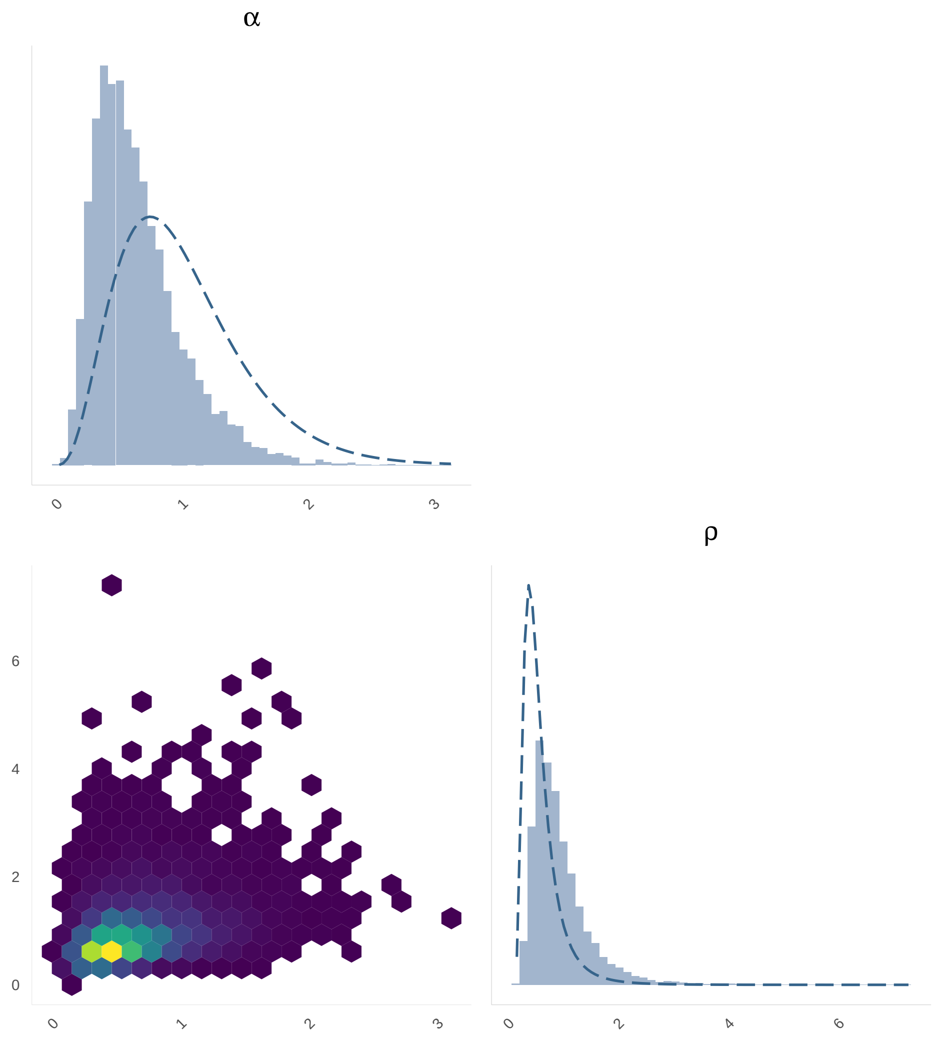


Figure S6: Marginal and joint distributions for hyperparameters $\rho$ and $\alpha$ for the analysis of *N. gonorrhoeae*.

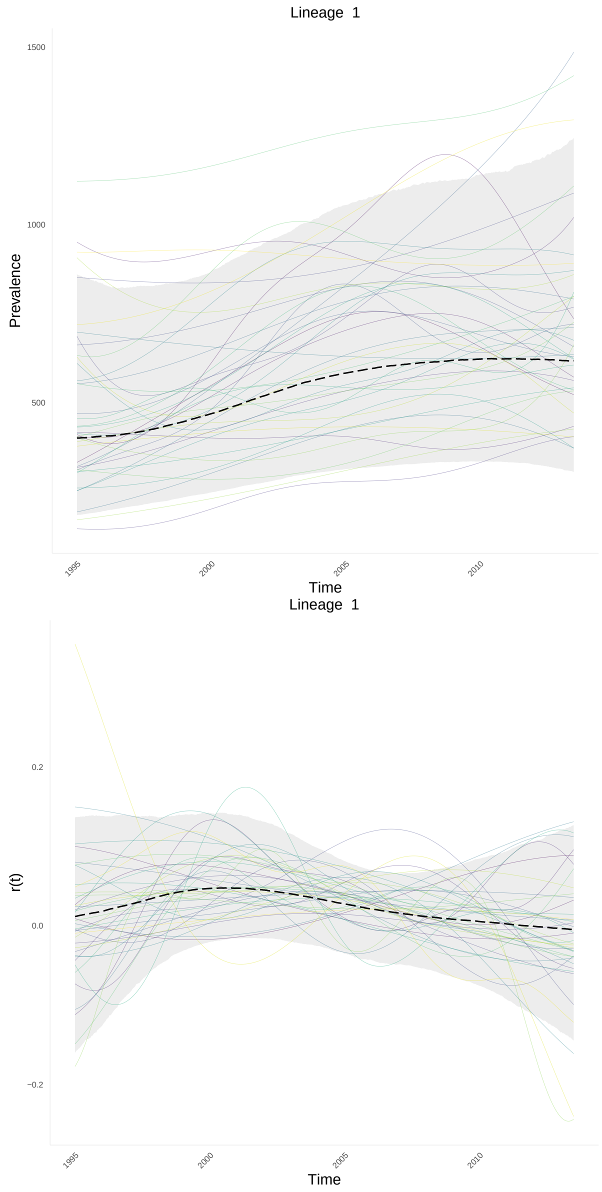


Figure S7: Estimated susceptible lineage dynamics for *N. gonorrhoeae*.

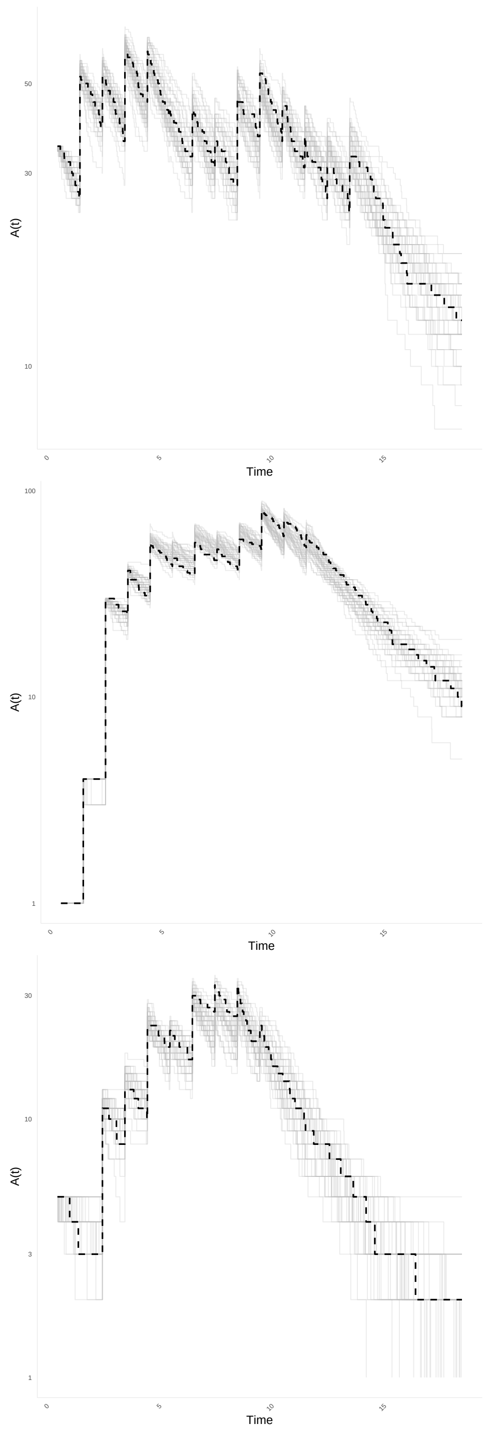


Figure S8: Posterior predictive trajectories of $A(t)$ for the susceptible lineage (top), resistant lineage 1 (middle) and resistant lineage 2 (bottom) of *N. gonorrhoeae*.
